# Spontaneous Dimerization and Distinct Packing Modes of Transmembrane Domains in Receptor Tyrosine Kinases

**DOI:** 10.1101/2024.05.09.593448

**Authors:** Lev Levintov, Biswajit Gorai, Harish Vashisth

## Abstract

The insulin receptor (IR) and the insulin-like growth factor-1 receptor (IGF1R) are homodimeric transmembrane glycoproteins that transduce signals across the membrane on binding of extracellular peptide ligands. The structures of IR/IGF1R fragments in apo and liganded states have revealed that the extracellular subunits of these receptors adopt Λ-shaped configurations to which are connected the intracellular tyrosine kinase (TK) domains. The binding of peptide ligands induces structural transitions in the extracellular subunits leading to potential dimerization of transmembrane domains (TMDs) and autophosphorylation in TKs. However, the activation mechanisms of IR/IGF1R, especially the role of TMDs in coordinating signal-inducing structural transitions, remain poorly understood, in part due to the lack of structures of full-length receptors in apo or liganded states. While atomistic simulations of IR/IGF1R TMDs showed that these domains can dimerize in single component membranes, spontaneous unbiased dimerization in a plasma membrane having physiologically representative lipid composition has not been observed. We address this limitation by employing coarsegrained (CG) molecular dynamics simulations to probe the dimerization propensity of IR/IGF1R TMDs. We observed that TMDs in both receptors spontaneously dimerized independent of their initial orientations in their dissociated states, signifying their natural propensity for dimerization. In the dimeric state, IR TMDs predominantly adopted X-shaped configurations with asymmetric helical packing and significant tilt relative to the membrane normal, while IGF1R TMDs adopted symmetric V-shaped or parallel configurations with either no tilt or a small tilt relative to the membrane normal. Our results suggest that IR/IGF1R TMDs spontaneously dimerize and adopt distinct dimerized configurations.

**TOC Graphic:** 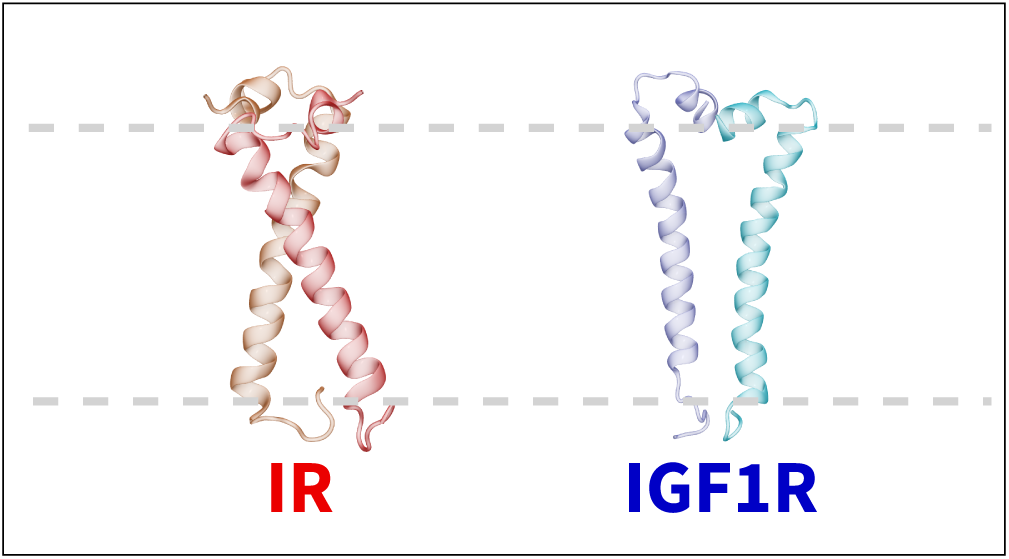

## Introduction

Membrane proteins mediate numerous cellular functions including signaling and transport.^1–4^ A fundamental structural element in these proteins is an *α*-helical transmembrane domain (TMD) which spans the hydrophobic core of the cell-membrane.^5,6^ TMDs contribute to the activation of membrane-spanning receptors by facilitating conformational transitions.^7,8^ Specifically, the dimerization of a pair of TMDs is an important step in initiating signaling via receptor tyrosine kinases (RTKs).^9,10^

Two key members of the RTK superfamily are the insulin receptor (IR) and the type 1 insulin-like growth factor receptor (IGF1R) that are homodimeric transmembrane glycoproteins.^10,11^ Each monomer in IR/IGF1R is comprised of an extracellular *α*-subunit and a membrane-spanning *β*-subunit containing a TMD flanked by juxtamembrane regions and connected to an intracellular tyrosine kinase (TK) domain.^7,10^ The binding of insulin or insulin-like growth factors to the extracellular subunits of IR/IGF1R results in autophosphorylation in the cytoplasmic TK domains and further downstream signaling. ^12,13^ In the absence of ligands, the extracellular subunits of IR and IGF1R adopt symmetric Λ-shaped configurations with spatially separated TMDs. ^14–17^ However, upon ligand binding, the extracellular subunits of IR and IGF1R transition to Γ- and T-shaped configurations with TMDs located near each other.^18–21^

Despite an abundance of structural data on IR and IGF1R,^15,16,20–34^ the role of dimerized TMDs in the activation of these receptors remains unclear due to the absence of a full-length receptor structure containing TMDs.^10,35^ However, the solution structure of an isolated TMD has revealed a well-defined *α*-helical shape with predominantly non-polar hydrophobic residues spanning the hydrophobic membrane layer (Figure 1).^7^ This structure indicates a kink at residues G960 and P961 in IR TMD which could be important for dimerization and/or receptor activation. ^7,10^ The IGF1R TMD also adopts a well-defined *α*-helical structure with non-polar hydrophobic residues constituting the helix and a kink formed at P941 (Figure 1).^36^ Both TMDs contain three positively-charged residues near the C-terminus (IR: R980, K981, and R982; IGF1R: R960, K961, and R962), which can engage in salt-bridging interactions with the negatively-charged lipid head groups of the inner membrane leaflet (Figure 1), thereby potentially anchoring the linkage motifs and further guiding the movement of the intracellular kinase domains. ^7^

**Figure 1:**
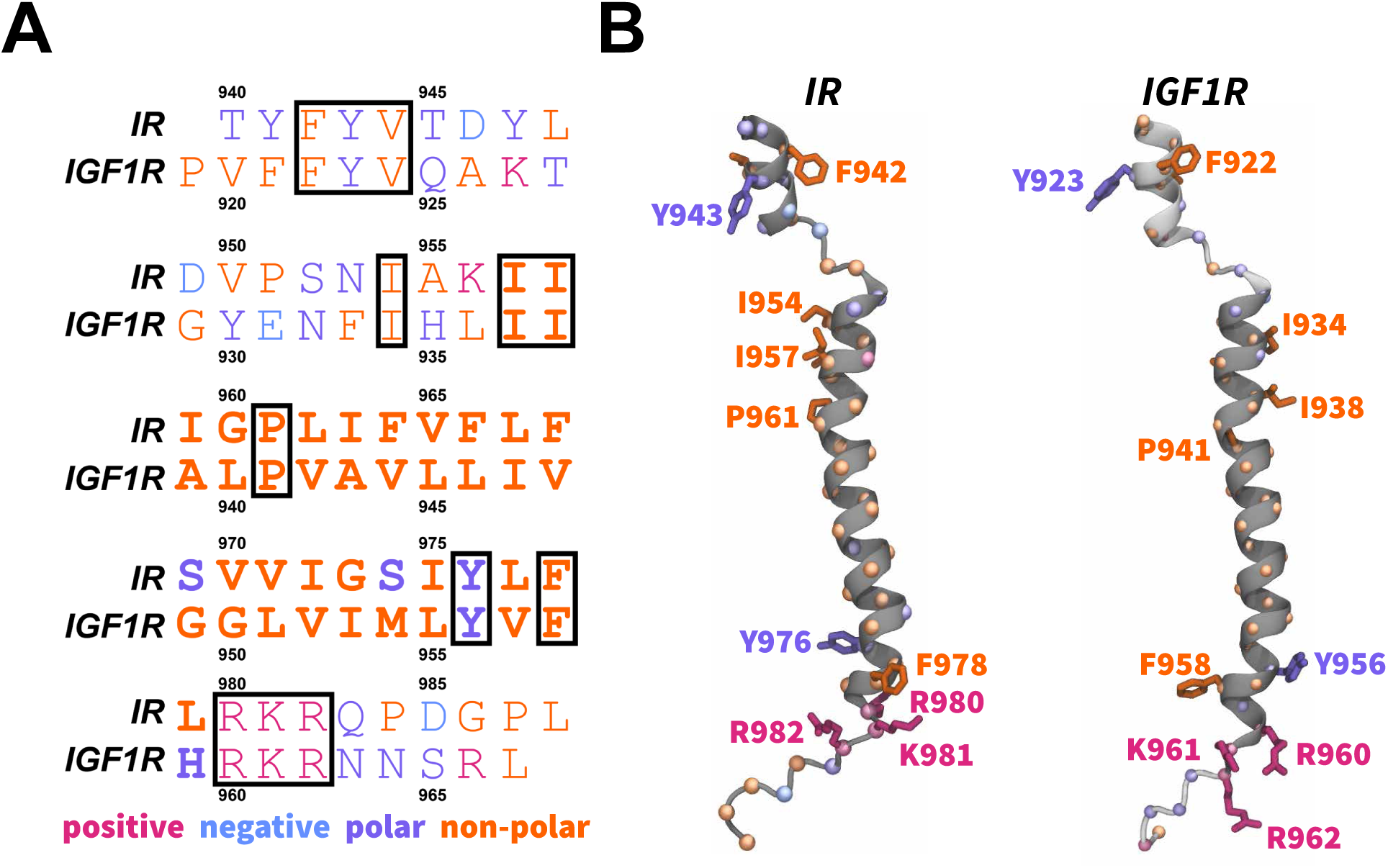
Sequences and structures of IR/IGF1R TMDs. (A) A sequence alignment of the IR and IGF1R TMDs with conserved residues enclosed in boxes. Bold letters signify residues embedded in the membrane and forming the transmembrane helix.^11^ (B) The structures of IR/IGF1R TMDs^7,37^ are shown in dark gray cartoons along with the modeled segments in light gray cartoons and with specific atoms/residues uniquely colored (spheres, C*_α_*; blue sticks, negatively-charged residues; magenta sticks, positively-charged residues; violet sticks, polar residues; and orange sticks, non-polar residues).

Initially, it was proposed that TMDs had a passive role in insulin signaling by simply anchoring the receptor to the membrane.^38^ However, further studies suggested that TMD-TMD interactions in the dimerized state also stabilized the active conformation of IR.^20,35^ It was also shown that substituting the TMD in IR for the TMD of glycophorin A inhibited insulin action.^39^ Additionally, mutations (IR: G960A, P961A, and V965D; IGF1R: V952E) in the IR/IGF1R TMDs or removal of TMDs were shown to affect downstream signaling and negative cooperativity in the receptors. ^40–43^ Several studies have also proposed that TMDs could dimerize in the inactive basal state of the receptor and dissociate upon ligand binding.^44–46^ Furthermore, a yo-yo model of receptor activation indicates that the kinase domains are released from an initially constrained position on ligand binding. ^47^ Thus, further understanding of the dynamics and interactions underlying the TMD dimerization is necessary to fully comprehend the receptor signaling mechanism.^35^

Molecular dynamics (MD) simulations have been proposed and used as a tool to validate or supplement structural data as well as to provide atomistic insights into the conformational dynamics of IR and IGF1R.^10,26,48–57^ MD simulations have also been used to characterize membrane proteins and their interactions with lipid molecules in the membrane.^58–61^ These studies have highlighted that the lipid composition modulates spatial configurations of transmembrane proteins and affects their dimerization process. MD simulations have also been utilized to characterize the orientation of the monomeric IR and IGF1R TMDs in a lipid bilayer.^36,48^ These atomistic simulations showed that the membrane-embedded residues of TMDs maintained the *α*-helical fold while exhibiting a tilt relative to the membrane. In one of these studies,^36^ MD simulations were conducted with IR and IGF1R TMD monomers embedded in several distinct membranes showing that the spatial orientation of TMDs is also influenced by the lipid composition, similar to other membrane proteins. ^60–62^ Furthermore, the dimerization process of IGF1R TMDs was probed using MD simulations which suggested that TMDs can form a dimer with interactions via a conserved proline residue.^22^

However, observing spontaneous TMD dimerization in a relatively larger membrane-protein system remains a challenging undertaking using atomistic MD simulations, even with modern supercomputing hardware.^63–65^ Therefore, various enhanced sampling techniques and special-purpose hardware have been applied to characterize the dimerization process of TMDs in membranes by more efficiently sampling conformational space of membrane-protein systems.^22,66–68^ These methods often require pre-defined collective variables and biasing forces to probe slower biophysical processes such as dimerization.^69,70^

A promising alternative approach is to coarse-grain (CG) a protein/membrane system by reducing the number of degrees of freedom while preserving the chemical properties of the system.^65,71,72^ This approach often enables simulations of larger biomolecular systems and captures processes occurring at longer timescales, which are usually inaccessible to all-atom MD simulations.^65,71–73^ Specifically, CG MD simulations have been applied to characterize the dimerization of TMDs in other proteins, further highlighting the importance of lipid composition and the ability of CG simulations to capture the complex behavior of dimerization in membranes.^62,74–76^

Therefore, we developed CG models of IR/IGF1R TMDs to probe their spontaneous dimerization in a plasma membrane representative of a physiologically relevant lipid composition.^71,77,78^ Since the orientations of TMDs relative to the membrane or to each other in the full-receptor context have never been experimentally resolved, we embedded TMDs in the membrane in several distinct orientations aiming to obtain a broader conformational mapping during their dimerization process. The dynamics of TMD dimerization were then investigated via a total of 300 *µ*s CG MD simulations. Briefly, we discovered that TMD molecules can spontaneously associate irrespective of their initial orientations or sequences, signifying that IR/IGF1R TMDs display a natural tendency to dimerize in the plasma membrane. Upon dimerization, IR TMDs preferentially adopted X-shaped configurations with a ∼30*^◦^* tilt relative to the membrane, while IGF1R TMDs preferentially adopted V-shaped or parallel configurations with no significant tilt relative to the membrane.

## Materials/Experimental Details

### Coarse-Grained Modeling

#### IR and IGF1R TMDs

We obtained the initial atomic coordinates for the human construct IR_940_*_−_*_988_ containing the TMD (hereafter referred to as IR TMD) from the first frame of the NMR structure (PDB code 2MFR).^7^ Furthermore, we modeled the tertiary structure of the human construct IGF1R_919_*_−_*_967_ containing TMD (hereafter referred to as IGF1R TMD) using the MODELLERv9.20.^79^ We used the structure of the IR TMD (PDB code 2MFR)^7^ as a template during model building using the homology modeling approach.^80^ We aligned the sequences for IR and IGF1R TMDs (Figure 1A) and generated 200 models of IGF1R-TMD using the MODELLER. We selected the best model based on the lowest discrete optimized protein energy (DOPE) score (Figure 1B).^81^ We further generated CG models for IR and IGF1R TMDs from the corresponding all-atom structures using the MARTINI force-field version 2.2 (Figure 2A).^71,77,82^

**Figure 2:**
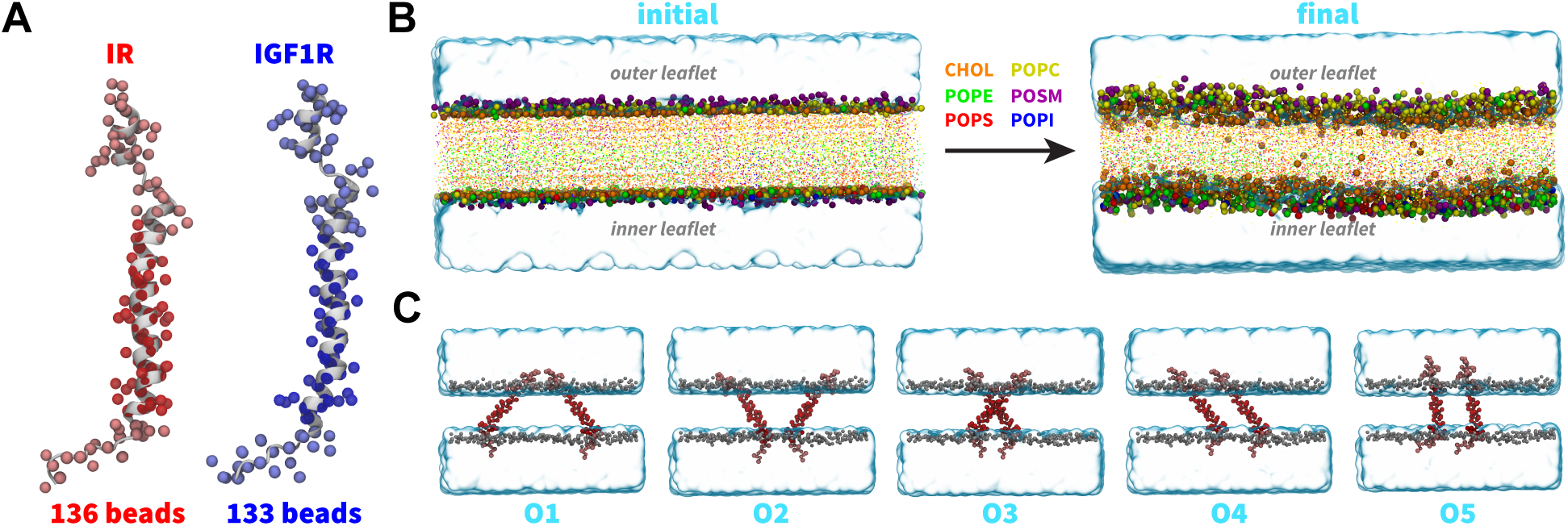
CG modeling and system setup. (A) CG models of IR TMDs (red spheres) and IGF1R TMDs (blue spheres) superimposed on the corresponding all-atom structures (white cartoons) with residues spanning the membrane highlighted in darker color spheres. (B) The initial CG lipid bilayer was subjected to equilibration and production simulations. Lipid head groups and tails are shown as uniquely colored spheres and points, respectively. Water molecules in the simulation domain are represented as a blue volumetric surface. (C) Side-view snapshots of simulation domains of CG IR TMDs embedded in a lipid bilayer in each distinct initial orientation (labeled O1 through O5). The setup for simulations of CG models of IGF1R TMDs was similar to those of IR TMDs.

#### Plasma membrane

We used the MARTINI builder in the CHARMM-GUI^83,84^ tool to construct a CG bilayer lipid membrane (20 nm × 20 nm × 4 nm) containing 1600 lipids (800 lipids per leaflet). The composition of the lipid bilayer was set to mimic the biological composition of a plasma membrane with the outer leaflet containing a mixture of 360 cholesterol molecules, 248 POPC lipids, 136 POSM lipids, and 56 POPE lipids while the inner leaflet contained 328 cholesterol molecules, 168 POPE lipids, 120 POPC lipids, 80 POPS lipids, 72 POSM lipids, and 32 POPI lipids. We solvated the lipid membrane with 15312 polarizable CG water molecules while keeping the membrane domain free of water molecules. We ionized the system with 314 Na^+^ and 202 Cl*^−^* ions at a salt concentration of 150 mM (Figure 2B).

After generating the solvated and ionized membrane system, we equilibrated it according to the following simulation protocol. In the first step, we performed an energy minimization for 5000 steps using the steepest-descent algorithm with a force tolerance of 100 kJ mol*^−^*^1^ nm*^−^*^1^. In the second step, we performed equilibration in the NVT ensemble using the Berendsen thermostat with a coupling time of 0.1 ps and temperature of 300 K for 10 ns followed by equilibration in the NPT ensemble using the Berendsen barostat at 1 atm pressure for 100 ns. During initial equilibration steps, we applied harmonic restraints (k = 1000 kJ mol*^−^*^1^ nm*^−^*^2^) on the polar beads in the lipid heads (namely ROH and PO4 beads of cholesterol and other lipids, respectively). In the last step, we performed unrestrained simulation in the NPT ensemble for 2 *µ*s with a timestep of 25 fs (Figure 2B). We maintained the temperature at 300 K and the pressure at 1 atm using the V-rescale thermostat and the Parrinello-Rahman barostat, respectively, in the unrestrained simulation.

#### System Setup and Simulation Details

To study the dimerization of TMDs within the plasma membrane, we generated separate systems with a pair of IR/IGF1R TMDs embedded in an equilibrated lipid bilayer in five distinct orientations, such that the N-terminus and the C-terminus of IR/IGF1R TMDs were directed toward the outer and inner leaflets, respectively (Figure 2C). In each system, IR/IGF1R TMDs were separated by a distance of 25 Å computed between the closest residues in a pair of TMDs. After embedding TMDs in the membrane, we deleted lipids located within 4 Å of these domains to remove any steric clashes. The resulting CG IR and IGF1R TMD systems contained ∼50000 and ∼58000 atoms, respectively (Table 1).

**Table 1:** Details on simulation systems. For each orientation of IR/IGF1R TMDs, listed are the system size, inter-center-of-mass distance (d_HH_) between a pair of TMDs, initial tilt (θ) and crossing (Ω) angles. The metrics for IGF1R TMDs are shown in bold.

| Orientation<br>IR/IGF1R | System Size<br>(atoms) | $d_{\text{HH}}$ (nm) | $\theta$ (°) | $\Omega$ (°) | Production<br>length ($\mu\text{s}$) |
| --- | --- | --- | --- | --- | --- |
| O1 | 50822 | 7.2 | 45 | 0 | 30 (10 $\mu\text{s}$ x 3) |
| **O1** | **58464** | **7.2** | **45** | **0** | **30 (10 $\mu\text{s}$ x 3)** |
| O2 | 51159 | 3.5 | 45 | 0 | 30 (10 $\mu\text{s}$ x 3) |
| **O2** | **58403** | **3.5** | **45** | **0** | **30 (10 $\mu\text{s}$ x 3)** |
| O3 | 50330 | 2.4 | 45 | 45 | 30 (10 $\mu\text{s}$ x 3) |
| **O3** | **58408** | **2.4** | **45** | **45** | **30 (10 $\mu\text{s}$ x 3)** |
| O4 | 50895 | 3.5 | 45 | 0 | 30 (10 $\mu\text{s}$ x 3) |
| **O4** | **58459** | **3.5** | **45** | **0** | **30 (10 $\mu\text{s}$ x 3)** |
| O5 | 48940 | 3.1 | 0 | 0 | 30 (10 $\mu\text{s}$ x 3) |
| **O5** | **58505** | **3.1** | **0** | **0** | **30 (10 $\mu\text{s}$ x 3)** |

For each system of IR/IGF1R TMDs, we conducted three independent 10-*µ*s-long production CG MD simulations (Table 1). Additionally, we conducted three independent 5-*µ*s-long CG MD simulations of the monomeric IR/IGF1R TMDs embedded in an equilibrated lipid bilayer in the O5 orientation. All CG MD simulations were conducted in the NPT ensemble with a 25 fs timestep. The coordinates from each simulation trajectory were saved at every 50 ps. The temperature and pressure were maintained at 300 K and 1 atm with V-rescale thermostat and Parrinello-Rahman barostat. A nonbonded cut-off of 11 Å was used for both Coulombic and van der Waals interactions. The periodic boundary conditions and semi-isotropic pressure coupling was used across all CG MD simulations. Beyond the cutoff for Coulombic interactions, the dielectric constant was set to 2.5. All simulations were conducted using the GROMACSv2020.4^85^ software package combined with the MARTINI force-field version 2.2 which resulted in the overall 300 *µ*s dataset (Table 1).^71,77,82^ The analyses of all trajectories were carried out using the tools in GROMACS^85^ and Visual Molecular Dynamics (VMD) software.^86^

### Conformational Metrics

#### Backmapping of CG models

We followed a procedure developed by Wassenaar *et al.*^87^ to convert representative structures from CG simulations into atomistic models to obtain additional insights about the atomic-scale processes from CG simulations.

#### Inter-helical distance (d_HH_)

We calculated the inter-helical distance between the centers-of-mass of two TMD helices (IR: residues 953 through 979; IGF1R: residues 937 through 959) across all CG MD simulations. According to this metric, the range of distances between dimerized IR/IGF1R TMDs in the X-shaped/V-shaped configurations was 1-1.3 nm and in the parallel configuration was 0.8-1 nm. Therefore, we defined the dimerized state when TMDs were located within 1.3 nm of each other.

#### Tilt (θ) and crossing (Ω) angles

We defined the tilt angle (θ) of each TMD helix (IR: residues 957 through 979; IGF1R: residues 937 through 959) with respect to the membrane by computing the angle between the vector projected along the TMD helical axis and the vector normal to the membrane surface. Furthermore, we defined the crossing angle (Ω) between each TMD pair by computing the angle between two vectors projected along the axis of each TMD helix. The tilt and crossing angles were calculated across all CG simulations initiated from five distinct TMD configurations and used to obtain the probability distributions of θ and Ω, respectively. In the monomeric IR/IGF1R TMD simulations, only the tilt angle analysis was performed.

#### Root mean squared fluctuation (RMSF)

We calculated the RMSF per residue based on the protein backbone atom (name BB) to characterize the flexibility of each TMD residue. The RMSF values were averaged over three independent CG simulations for simulations initiated from each initial orientation.

#### Free energy calculation

We estimated the free energy change along θ following the histogram method previously used to characterize the dimerization process of transmembrane proteins.^62,88^ The free energy estimate is given by *U* = −*k_B_T ln*[*P* (θ)] where *k_B_* is Boltzmann’s constant, *T* is the temperature, and *P* (θ) is the probability of observing a value of θ. The free energy estimates were averaged over three CG simulations for each receptor orientation. This procedure was repeated for calculating the free energy change along Ω. **Cluster analysis:** We performed the cluster analysis using the GROMACSv2020.4^85^ software following the clustering algorithm by Daura et al.^89^ We extracted dimerized states from all CG simulations and clustered them based on similarity using the RMSD cutoff of 0.9 nm.

#### Dimerization interface analysis

We analyzed the dimerization interface by computing averaged distances between the centers-of-mass of each residue pair in the TMD helices. Based on the inter-helical distance analysis, the coordinates of dimerized TMD configurations were saved.

## Results

### Spontaneous Dimerization of IR/IGF1R TMDs

To study the dimerization process of TMDs, we inserted a pair of IR or IGF1R TMDs in an equilibrated CG model of the plasma membrane in five distinct orientations (Figure 2C; Table 1). Three independent CG simulations (each 10 *µ*s long) were conducted for each initial orientation of a pair of TMDs (Table 1). We used the inter-center-of-mass distance between a pair of TMD helices as a metric to monitor the formation of a TMD dimer (d_HH_; Figure 3). An increase in the d_HH_ value signifies that TMDs diffused away from each other while a decrease signifies that TMDs moved toward each other. Across all CG simulations, we observed that each TMD initially diffused in the membrane plane with the diffusive search leading to an encounter with the other TMD to form a dimer (Figure 3). Despite the initial diffusion of IR/IGF1R TMD monomers away from each other (d_HH_ up to 8 nm) across several simulations, these molecules could still diffuse closer and spontaneously dimerize under 10 *µ*s timescale (Figure 3). Upon dimerization, TMD molecules maintained a stable dimeric state for the remainder of each simulation without any transient dissociation (Figure 3). Thus, independent of their initial orientations, spontaneous dimerization of TMDs was observed in all simulations within the 10 *µ*s timescale.

**Figure 3:**
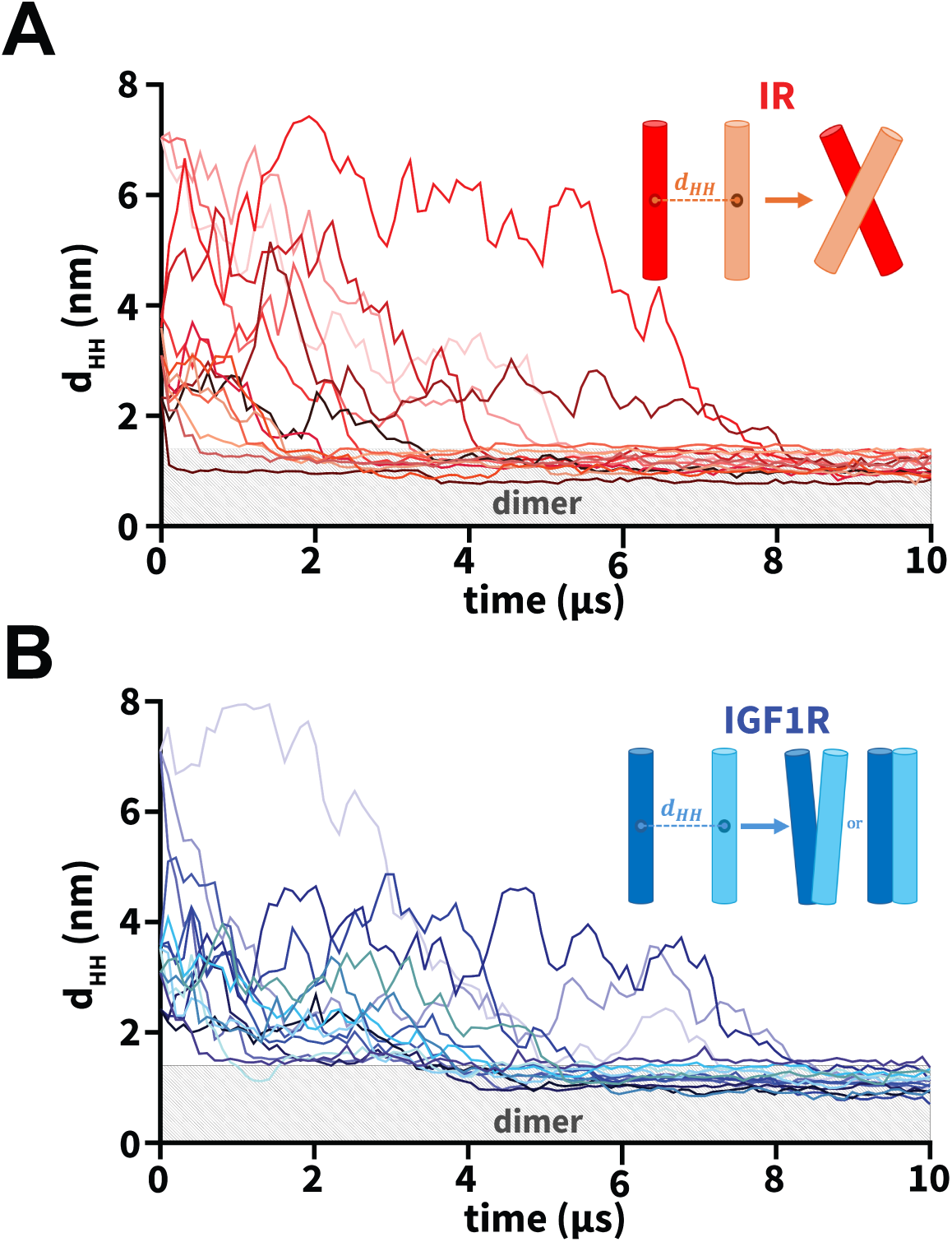
Dimerization of TMDs. (A and B) The uniquely-colored traces of the interhelical distances (d_HH_) vs. simulation time highlight the dimerization of IR/IGF1R TMDs for each CG simulation. The light gray rectangle in each plot marks distances below 1.3 nm indicating dimerized configurations. Insets in each plot highlight a schematic representation of the dominant dimerized state of each pair of IR/IGF1R TMDs: X-shaped (IR) and V-shaped or parallel (IGF1R).

Furthermore, we measured the interfacial buried surface area (BSA) between a pair of TMD molecules to assess the interaction interface between TMDs in dimerized configurations (Figure S1). We observed an increase in BSA from the initial value of zero when the interhelical distance between TMDs was in the range of 3 nm to 4 nm (Figure S1). The BSA increase was due to the interactions formed between the N-termini of TMDs which tend to initiate TMD association across various IR and IGF1R simulations (Figure S2). As the interhelical distance decreased to values below 1.3 nm, BSA increased to values ranging between 8 nm^2^ and 24 nm^2^, signifying the association of TMD molecules (Figure S1). Upon dimerization, IR and IGF1R TMDs displayed distinct modes of helical packing, with IR TMDs predominantly forming X-shaped configurations and IGF1R TMDs forming either V-shaped or parallel configurations (inset; Figure 3).

Additionally, we computed the RMSF per residue to assess the flexibilities of TMD residues across all CG simulations (Figure S3). We observed that residues embedded in the membrane (IR: I953 through L979; IGF1R: I937 through H959) were less flexible in comparison to residues in the rest of the structure (Figure S3). These observations are consistent with prior atomistic simulations of IR TMDs which showed decreased RMSF values for residues embedded in the membrane in comparison to the solvent-exposed residues, signifying that the CG models capture the conformational behavior observed in atomistic simulations of IR/IGF1R TMDs.^37^ Overall, CG simulations demonstrated spontaneous self-association of IR/IGF1R TMDs independent of their initial orientations or differences in their sequences.

### Packing Modes of IR/IGF1R TMDs

We characterized the dimerization interfaces by identifying dominant dimerized configurations via an RMSD-based clustering analysis (Figures 4, S4).^89^ In simulations of IR TMDs, two largest conformational clusters comprising ∼80.8% of all sampled conformations, displayed the formation of X-shaped configurations (Figure 4A). A key difference between the X-shaped configurations was the orientation of the N-terminal motifs (residues 940 through 953) which influenced the location of the intersection point between two TMDs, thereby inducing tilted configurations of TMDs relative to the membrane and to each other (Figure 4A). In both clusters, the intersection point between IR TMDs was located near the kink formed by G960 and P961 residues (Figure 4A). Additionally, non-polar residues from each TMD helix engaged in inter-helical hydrophobic interactions, namely through the residues G960, P961, F964, F966, and F968 (Figure 4A). For IGF1R TMDs, we observed the formation of either a V-shaped configuration (C1; Figure 4B) or a tightly packed parallel configuration (C2; Figure 4B) which were distinct from the X-shaped configurations adopted by a pair of dimerized IR TMDs. The V-shaped configuration also exhibited parallel helical packing with a kink in TMDs near the P941 residue but the N-terminal motifs were intercalated between the TMD helices, thereby rotating residues N932 through I947 away from each other (C1; Figure 4B). Overall, hydrophobic interactions defined the TMD-TMD interface in the dimerized states irrespective of the receptor type or initial configuration.

**Figure 4:**
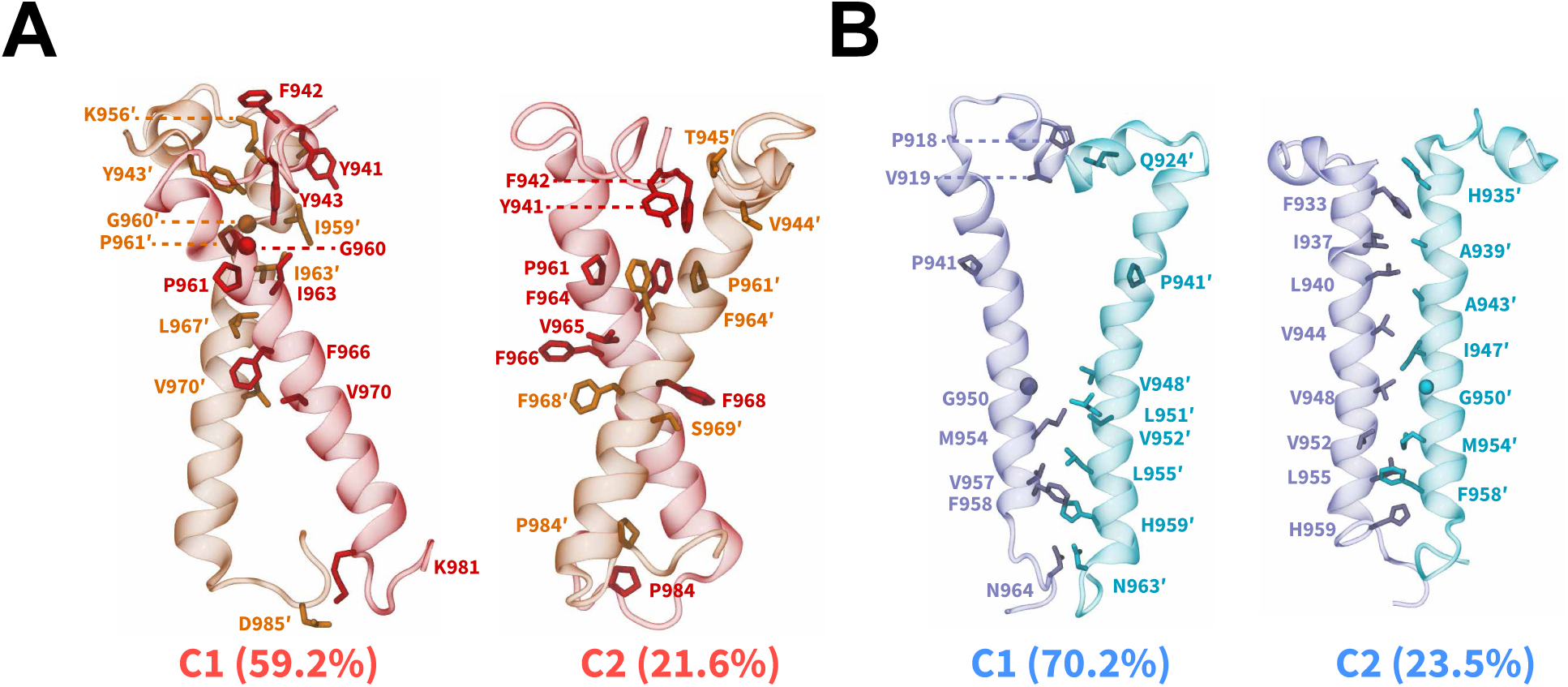
Dimerized configurations and packing modes of IR/IGF1R TMDs. Sideview snapshots of the averaged structures (cartoon) from two largest clusters of the dimerized states derived from CG simulations of (A) IR and (B) IGF1R TMDs. The residues in the interface are labeled and shown as sticks except G960 (IR) and G950 (IGF1R) which are shown as spheres. The names of conformational clusters (C1 and C2) and their sizes (%) are labeled. See also Figure S4.

Additionally, the clustering analysis showed the formation of several smaller-sized clusters (Figure S4). Specifically, IR TMDs adopted parallel configurations which in total constituted ∼17.6% of dimerized configurations (C3 and C4; Figure S4A). These parallel configurations differed from each other by their relative orientation with respect to the membrane. However, these clusters likely represent intermediate dimerized configurations of IR TMDs, given the relatively smaller sizes of these clusters. A small cluster having 6.3% of total configurations was also observed in simulations of IGF1R TMDs, highlighting another parallel configuration (C3; Figure S4B). A key feature of this cluster was the presence of interactions between the N-terminal motifs of IGF1R TMDs which were absent in the C2 cluster.

Furthermore, we characterized the interfaces formed upon dimerization using the residue-contact-map analysis based on distances between the residue pairs (Figure S5). We observed that IR TMD dimers showed a wider distribution of residue pairs in close contact with each other, as indicated by their off-diagonal placement in the contact maps, also signifying the asymmetric helix-helix interface in IR TMDs (Figure S5A). On the contrary, we observed that IGF1R TMD dimers predominantly exhibited residue pairs at or near the diagonal in the contact maps, corresponding to a symmetric helix-helix interface (Figure S5B). Furthermore, TMDs for both IR and IGF1R showed tighter residue contacts near the N-termini (Figure S5B), suggesting that residues from the N-termini assist in stabilizing TMDs in the dimerized configurations.

### Relative Orientations of TMDs and Energetics of Dimerization

We further assessed the spatial orientations of TMD monomers and dimers relative to the membrane normal using a commonly defined tilt angle (θ) for transmembrane proteins (Figure 5A).^37,48,62^ In simulations of IR and IGF1R TMD monomers, we observed the θ distributions to be in the range of values between 0*^◦^* and 45*^◦^*, and between 0*^◦^* and 30*^◦^*, respectively (Figure S6A). Furthermore, the θ distributions for IR TMD dimers (Figures 5B, S6B) showed the range of values between 0*^◦^* and 50*^◦^*. The range of θ values observed in our simulations of IR TMD monomer and dimer were similar to each other and to the previously reported tilt angle values (between 0*^◦^* and 50*^◦^*) computed based on atomistic MD simulations of IR TMD monomers, thereby in agreement with our results. ^36,37,48^ However, for IGF1R TMD in both monomeric and dimeric states, we observed that the θ values ranged between 0*^◦^* and 30*^◦^* (Figures 5B, S6A, C), indicating a lower tilt (relative to the membrane normal) of IGF1R TMDs in comparison to IR TMDs. Prior simulation work of the monomeric IGF1R TMD reported the tilt angle to be in the range of 10*^◦^* to 40*^◦^*,^48^ similar to the values observed in our work. The distributions of θ as a function of d_HH_ (Figure S7) indicated that even upon dimerization (d_HH_ < 1.3 nm), IR/IGF1R TMDs adopted configurations with tilt angles similar to their dissociated (d_HH_ > 4 nm) and monomeric states (Figure S6A). Thus, dimerization of TMDs did not significantly constrain their propensities to tilt relative to the membrane (Figures S6A, S7).

**Figure 5:**
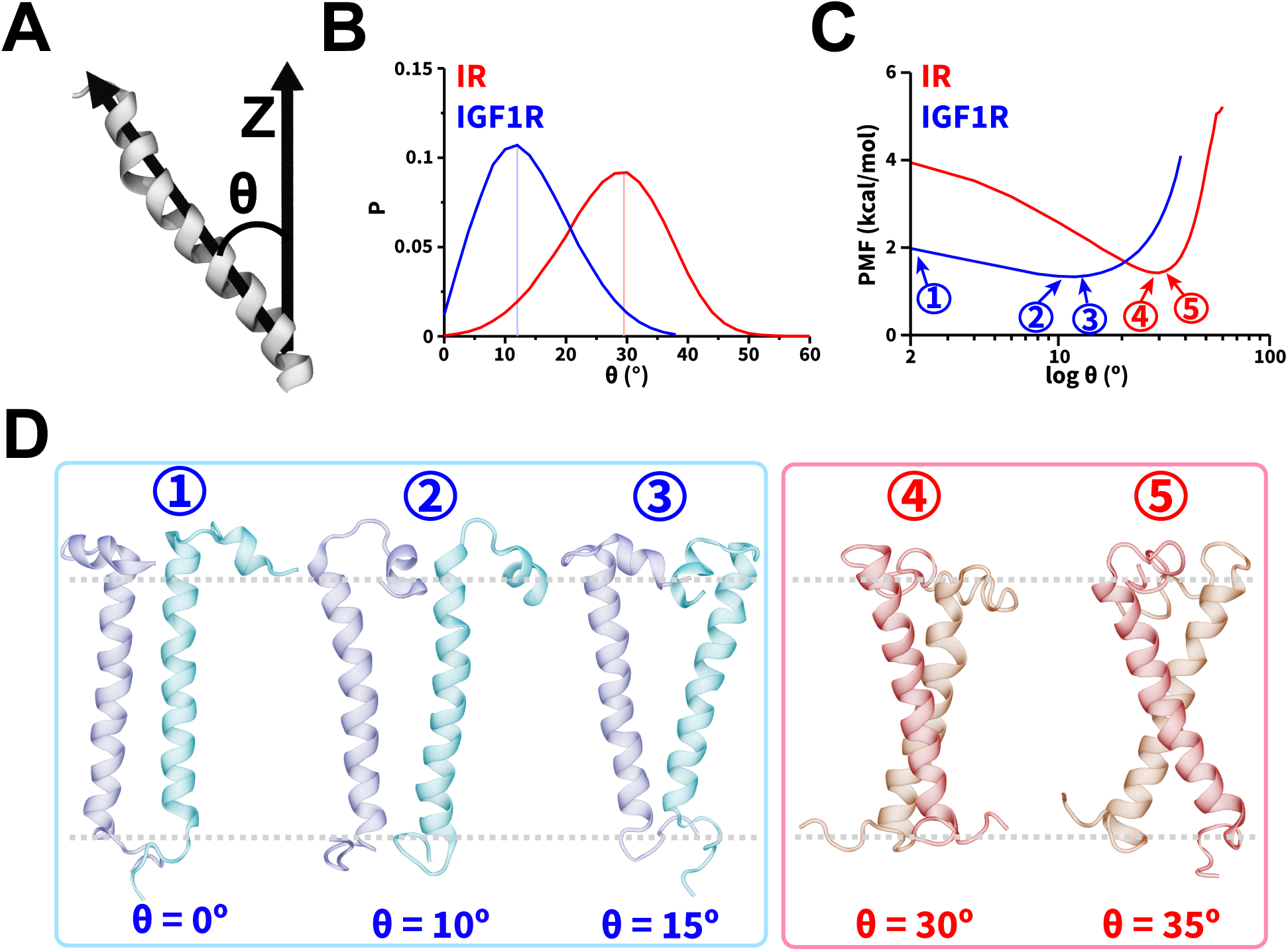
Orientations and energetics of dimerized TMDs. (A) A schematic highlighting the vectors used in defining θ, (B) probability distributions of θ, and (C) the potential of mean force (PMF) highlighting free energy change (kcal/mol) as a function of θ. In panels B and C, the traces from CG simulations of IR and IGF1R TMDs are shown in red and blue, respectively. (D) Snapshots showing dimerized TMD configurations corresponding to various values of θ labeled 1 through 5 in panel C with gray dashed lines highlighting the approximate thickness of the membrane. See also Figure S6.

We used the probability distributions of θ from simulations of the dimeric state to estimate the free energy change of dimerization of IR/IGF1R TMDs. This metric has been previously utilized in CG simulations to quantify the free energy change of transmembrane protein dimerization and showed good agreement with all-atom models. ^62,88^ We observed lower free energy values for 20*^◦^* < θ < 40*^◦^* with the lowest free energy value corresponding to a free energy minimum at θ = 30*^◦^* (Figure 5C). At this free energy minimum, IR TMDs adopted various X-shaped configurations with slightly altered θ values (Figure 5D). However, for IGF1R TMDs, lower free energy values were observed between 0*^◦^* and 20*^◦^* without a prominent free energy minimum (Figure 5C). The IGF1R TMD helices adopted either a parallel configuration (label 1; Figure 5D) or V-shaped configurations (labels 2 and 3; Figure 5D). Thus, IGF1R TMDs could adopt lower free energy configurations having 0*^◦^* < θ < 20*^◦^* (Figure 5C).

In addition to assessing the orientations of TMDs relative to the membrane, we computed the crossing angle (Ω) to characterize the relative orientation of TMDs with respect to each other in the dimerized states (Figure S8A). This is also a commonly used metric for characterizing helical packing in membrane-protein simulations.^74,75,90^ We observed that Ω fluctuated between 0*^◦^* and 35*^◦^* (IR TMDs) and between 0*^◦^* and 15*^◦^* (IGF1R TMDs) in all CG simulations (Figure S8B). Thus, Ω data showed that IR/IGF1R TMDs adopted either a parallel (Ω = 0*^◦^*) or slightly tilted (0*^◦^* < Ω < 15*^◦^*) configurations relative to each other (Figure S8B). However, IR TMDs further adopted configurations with Ω > 15*^◦^* which resulted in X-shaped configurations (Figures 4A, 5D). Thus, dimerized IR TMDs adopted conformations which were more inclined relative to each other than the helices in the dimerized IGF1R TMD configurations.

The free energy profiles as a function of Ω showed that in IR TMDs, Ω ranged between 0*^◦^* and 30*^◦^* with the free energy profile having no significant free energy minimum (Figure S8C). The shapes of the free energy profiles as a function of θ and Ω signify that IR TMDs favored tilted configurations relative to the membrane while predominantly adopting X-shaped configurations (Ω > 0*^◦^*) or forming parallel configurations relative to each other (Ω = 0*^◦^*) (Figure S8C). For IGF1R TMDs, we observed Ω ranging between 0*^◦^* and 15*^◦^* with a free energy minimum near Ω = 0*^◦^* (Figure S8C). Thus, IGF1R TMDs favored a parallel configuration relative to each other while adopting either a perpendicular (θ = 0*^◦^*) or slightly tilted V-shaped configurations (Figure 5D) relative to the membrane normal.

The distributions of Ω (Figure S9) showed that as IR TMDs approached each other, the range of values for Ω increased in comparison to the values at greater separation distances, signifying conformational rearrangements during the formation of the X-shaped configurations (Figures 4A, 5D). However, the dimerization of IGF1R TMDs either did not alter the distributions of Ω or slightly increased Ω by ∼10*^◦^*, signifying a higher probability of parallel helical packing in comparison to IR TMDs (Figure S9). Thus, IR TMDs were more dynamic upon dimerization with a higher propensity to form X-shaped configurations while IGF1R TMDs were less tilted relative to each other, thereby forming V-shaped or parallel configurations.

## Discussion

Elucidating the role of TMD dimerization is a crucial step in understanding the mechanisms of activation of IR and IGF1R.^10^ It remains poorly understood if the initial Λ-shaped apo configuration of the extracellular IR domain is a crystallographic artifact resulting from the absence of TMDs or there are other contributing factors.^10^ In this work, we conducted CG simulations of a pair of IR and IGF1R TMDs to probe the spontaneous dimerization process in a membrane having a lipid composition representative of the plasma membrane. Importantly, the lipid composition of the plasma membrane has been shown to influence the conformations and orientations of various transmembrane proteins,^62,75^ including monomeric and dimeric IR/IGF1R TMDs through the application of all-atom and CG MD simulations.^36,48^ Therefore, we designed CG membrane models with the lipid composition of the plasma membrane,^91^ which has not been used in previous simulation studies of IR/IGF1R TMDs.^22,36,37,48^ The thickness of our CG membrane was ∼40 Å, which embodied the entire hydrophobic transmembrane regions (∼36 Å) of the utilized IR/IGF1R TMD fragments.

To observe unbiased and spontaneous dimerization of TMDs, we initiated long timescale (10 *µ*s) CG MD simulations with multiple independent initial orientations in dissociated configurations of TMDs. On conducting CG MD simulations of both IR/IGF1R TMDs, we observed their spontaneous and unbiased dimerization independent of their initial orientations. Upon dimerization, IR/IGF1R TMDs stably maintained their associated states. Thus, CG simulations showed that TMD molecules can spontaneously dimerize without any external bias, indicating their natural propensity for self-association.

The residue-fluctuation RMSF analysis showed that the helical motifs of TMDs (IR: I953 through L979; IGF1R: I937 through H959) were more rigid than the residues in the termini that are exposed to the solvent, as also suggested in prior all-atom simulations of IR TMDs.^37^ Furthermore, the flexible residues from the N-termini were observed to facilitate dimerization across all systems, signifying their potential importance in the dimerization process, especially in bringing the type-III fibronectin (FnIII-3) domains closer to each other. Upon dimerization, the N-terminal residues from the opposite TMDs continued interacting with each other, contributing to the stability of the dimeric configuration and potentially stabilizing the overall extracellular domains of IR/IGF1R.

We also calculated the tilt and crossing angles (θ and Ω) to characterize the spatial orientations of TMDs relative to the membrane normal and to each other, respectively. Based on the tilt angle analysis, we observed that both IR and IGF1R TMDs were tilted relative to the membrane normal while IR TMDs adopted configurations with higher tilt values than IGF1R TMDs. Specifically, the free energy profiles showed that the tilt angle was confined between 20*^◦^* and 40*^◦^* for IR TMDs with the most favorable configuration at θ∼30*^◦^*. IGF1R TMDs were tilted between 0*^◦^* and 20*^◦^* without any significant free energy minimum. Several prior structural studies^10,20,21^ have proposed that IR TMDs might occupy tilted configurations upon dimerization which is in agreement with our results. However, the relatively low resolution of cryo-EM maps in the transmembrane region of IR hinders the accurate modeling of this motif.^10,20,21^ Additionally, prior simulation work ^36,48^ of monomeric IR/IGF1R TMDs indicated that these TMDs could adopt configurations with a wide range of tilt angles confined between 0*^◦^* and 50*^◦^*. We also observed the propensity of dissociated TMDs (d_HH_ > 4 nm) as well as of monomeric TMDs to adopt a broad range of tilted configurations (IR: 0*^◦^* to 50*^◦^*; IGF1R: 0*^◦^* to 30*^◦^*), thereby in agreement with prior computational work of monomeric TMDs (IR: 0*^◦^* to 50*^◦^*; IGF1R: 10*^◦^* to 40*^◦^*).^36,37,48^

The crossing angle (Ω) analysis demonstrated that IR TMDs were prone to adopting various X-shaped configurations with a broad distribution of Ω values. Furthermore, these configurations were preferentially adopting a tilted configuration relative to the membrane normal according to the free energy analysis. IGF1R TMDs on the contrary favored V-shaped and parallel configurations while either adopting perpendicular configurations or slightly tilted configurations relative to the membrane (Ω < 15*^◦^*). Overall, the analysis of θ and Ω suggested that IR TMDs were more dynamic than IGF1R TMDs with a broader range of possible θ and Ω angles. IR TMDs adopted X-shaped configurations in the dimerized state while IGF1R TMDs adopted V-shaped or parallel configurations.

Currently, no structural data are available for the dimerized states of IR or IGF1R TMDs. Prior NMR work suggested that IR TMDs in their oligomeric forms in micelles could adopt various configurations with different interfaces.^7,37^ In our work, the conformational clustering analysis revealed that dimerized IR TMDs predominantly adopted X-shaped configurations. The interfacial residue contact maps further showed that IR TMDs predominantly formed asymmetric helix-helix interfaces, signifying that IR TMDs altered their orientations relative to each other prior to adopting an optimal dimeric configuration with the N-terminal residues engaged in a lateral helical packing mode. Additionally, we observed hydrophobic interactions among residues G960, P961, F964, F966, and F968 in the helix-helix interface of dimerized IR TMD configurations. These residues are located at or near the kink formed by the G960 and P961 residues in IR TMD helices, which is a common structural feature across various transmembrane proteins.^92–94^ The IR TMDs exhibited an *α*-helical shape in the X-shaped configuration which was in agreement with prior computational data which showed that IR TMDs maintained their *α*-helical folds throughout all-atom MD simulations. ^36,48^

Previous structural studies of a monomeric IR TMD in micelles highlighted the presence of a kink formed at G960/P961 residues.^7,10^ Furthermore, G960A and P961A mutations in IR TMDs resulted in an altered helical configuration of a monomeric IR TMD in micelles. ^42^ Based on these studies, it has been proposed that a kink at G960/P961 residues could increase the flexibility of the IR TMD, thus allowing it to alter its configuration for optimal dimerization. Our observation of the X-shaped configuration of dimerized IR TMDs with an intersection point near the G960/P961 kink further suggests that these residues could act as pivot points for TMD molecules to rotate into an optimal configuration, thereby in agreement with prior experimental work. ^7,10,42^ Based on the cryo-EM structures of the extracellular IR region it was proposed that the fibronectin domains FnIII-2/FnIII-2*^′^* and FnIII-3/FnIII-3*^′^* were conformationally flexible, thereby allowing them to switch between distinct states.^10,18–21^ To undergo these transitions, the linkages between FnIII-3 domains and TMDs should also be flexible. In the X-shaped configuration observed in our simulations, flexible linkages between the TMDs and the extracellular domains of IR would not sterically block each other and could potentially bring the FnIII-3 domains closer to each other, as observed in various experimental structures. ^18–21^ In several cryo-EM structures,^21,25^ the C-terminal (membrane-proximal) residues from two FnIII-3 domains are separated by ∼15-18 Å which is close to the values observed in the X-shaped configurations of IR TMDs (C1: 11.5 Å; C2: 13.6 Å; Fig. 4). Furthermore, the crystal structure of the IR TK domains which included additional juxtamembrane (intracellular) residues showed that the juxtamembrane motifs were oriented in a trans configuration across the TK domains. ^10,95^ Our conformational clustering analysis of IR TMD simulations also demonstrated the formation of the X-shaped configuration with the juxtamembrane residues located across from each other.

IGF1R TMDs adopted different dimerized configurations in comparison to IR TMDs, forming a more symmetric helix-helix interface with either a V-shaped configuration or a tightly packed parallel configuration. Thus, the dimerization mechanism of IGF1R TMDs could be different from the dimerization mechanism of IR TMDs. Specifically, the kink at P941 was preserved in the V-shaped dimerized configuration of IGF1R TMDs. This kink oriented the helical segment of IGF1R TMDs away from each other, which generated a bent configuration and provided additional space for the N-terminal motifs to intercalate between the TMD helices. Previous biochemical studies of a monomeric IGF1R TMD reported the formation of a bent structure near the kink, which is consistent with our observation. ^22,36^ Furthermore, while IGF1R TMDs were not as tilted relative to the membrane and to each other as IR TMDs, the N-termini in IGF1R TMDs were not sterically overlapping with each other. Thus, due to the structural kink introduced by the P941 residue, the N-terminal motifs adopted spatially closer configurations which could potentially induce conformational rearrangements in the FnIII-3 domains, bringing them to a more compact configuration and assisting in the transition of IGF1R into the activated state. The distance between the C-terminal residues (membrane-proximal) of FnIII-3 domains in one of the cryo-EM structures^96^ of IGF1R is reported to be ∼12 Å, which is close to the distance value between the N-terminal residues (extracellular) in the C1 configuration of IGF1R TMDs observed in our work (14.4 Å). Additionally, in another cryo-EM structure of IGF1R,^19^ the distance between the C-terminal residues of FnIII-3 domains is ∼27 Å which is close to the distance value between the N-terminal residues in the C2 configuration of IGF1R TMDs (22.2 Å). Thus, IGF1R TMD configurations observed in our simulations are potentially capturing the experimentally observed conformational behavior of IGF1R.

Overall, IR and IGF1R TMDs showed natural propensities of dimerization into distinct configurations which could potentially stabilize ligand-bound receptor configurations or facilitate conformational transitions in IR/IGF1R from their inactive states into active states. The kinks at G960/P961 (IR) and P941 (IGF1R) residues could further assist these conformational transitions by providing structural flexibility to alter the orientations of TMDs and bringing the FnIII-3 domains closer. Therefore, our work showed that CG modeling is a useful tool to study a complex biophysical process involving dimerization of transmembrane domains, which provides enhanced insights into their role in the activation of tyrosine kinase receptors of the insulin family.

## Conclusion

The role of TMD dimerization in the signal transduction mechanism of IR and IGF1R remains poorly understood, mainly due to the lack of structural details of the full-length receptors. In this work, we used CG MD simulations to probe the dimerization process of IR/IGF1R TMDs in a plasma membrane representative of physiologically relevant lipid composition. Since the initial orientation of TMDs relative to the plasma membrane in the context of the full-length receptor is not known, we embedded these TMD molecules in the membrane in several distinct orientations, aiming to broaden the conformational mapping of TMD dimerization. We observed spontaneous dimerization of TMDs independent of their initial orientations and the TMD sequences without any transient dissociation, signifying that IR/IGF1R TMDs are susceptible to forming a dimerized configuration even in the absence of the extracellular receptor domain. Furthermore, IR/IGF1R TMD association was facilitated by the N-terminal residues, potentially signifying their important role in bringing the FnIII-3 domains from the extracellular fragment of IR/IGF1R toward each other. TMD spatial orientation analysis revealed that both IR and IGF1R TMDs remained tilted relative to the membrane normal with IR TMDs being more tilted in comparison to IGF1R TMDs. Upon dimerization, IR TMDs predominantly adopted X-shaped configurations, while IGF1R TMDs predominantly adopted V-shaped or parallel configurations with a small tilt relative to the membrane. Both of these configurations preserved the kinks at G960/P961 (IR) and P941 (IGF1R) residues which contributed to the formation of distinct dimerized TMD configurations. These dimeric configurations of TMDs could potentially stabilize ligand-bound receptors and further assist in transitioning to their active states.

## Notes

The authors declare no competing financial interest.

## Acknowledgement

We acknowledge financial support from the National Institutes of Health (R35GM138217). We also acknowledge computational support through the following resources: Premise, a central shared HPC cluster at UNH supported by the Research Computing Center; BioMade, a heterogeneous CPU/GPU cluster supported by the NSF EPSCoR award (OIA-1757371).

## Supporting Information Available

The following data are presented in the supporting information.

- Additional BSA data, RMSF data, clustering analysis data, contact map data, tilt angle data, and crossing angle data from all CG simulations, structural insights from CG simulations on interactions between the N-termini of TMDs.

## References

(1) Liu, J.; Rost, B. Comparing function and structure between entire proteomes. Protein Sci. 2001, 10, 1970–1979.

(2) Nilsson, J.; Persson, B.; von Heijne, G. Comparative analysis of amino acid distributions in integral membrane proteins from 107 genomes. Proteins 2005, 60, 606–616.

(3) Cournia, Z. et al. Membrane protein structure, function, and dynamics: a perspective from experiments and theory. J. Membr. Biol. 2015, 248, 611–640.

(4) Jelokhani-Niaraki, M. Membrane proteins: structure, function and motion. Int. J. Mol. Sci. 2022, 24, 468.

(5) Killian, J. A.; von Heijne, G. How proteins adapt to a membrane–water interface. Trends Biochem. Sci. 2000, 25, 429–434.

(6) Sharpe, H. J.; Stevens, T. J.; Munro, S. A comprehensive comparison of transmembrane domains reveals organelle-specific properties. Cell 2010, 142, 158–169.

(7) Li, Q.; Wong, Y. L.; Kang, C. Solution structure of the transmembrane domain of the insulin receptor in detergent micelles. Biochim. Biophys. Acta - Biomembr. 2014, 1838, 1313–1321.

(8) Zhou, Y.; Wang, B.; Yuan, F. The role of transmembrane proteins in plant growth, development, and stress responses. Int. J. Mol. Sci. 2022, 23, 13627.

(9) Hubbard, S. R.; Miller, W. T. Receptor tyrosine kinases: mechanisms of activation and signaling. Curr. Opin. Cell. Biol. 2007, 19, 117–123.

(10) Lawrence, M. C. Understanding insulin and its receptor from their three-dimensional structures. Mol. Metab. 2021, 52, 101255.

(11) Adams, T.; Epa, V.; Garrett, T.; Ward, C. Structure and function of the type 1 insulin-like growth factor receptor. Cell. Mol. Life Sci. 2000, 57, 1050–1093.

(12) De Meyts, P. Insulin and its receptor: structure, function and evolution. Bioessays 2004, 26, 1351–1362.

(13) White, M. F.; Kahn, C. R. Insulin action at a molecular level–100 years of progress. Mol. Metab. 2021, 52, 101304.

(14) McKern, N. M. et al. Structure of the insulin receptor ectodomain reveals a folded-over conformation. Nature 2006, 443, 218–221.

(15) Croll, T. I.; Smith, B. J.; Margetts, M. B.; Whittaker, J.; Weiss, M. A.; Ward, C. W.; Lawrence, M. C. Higher-resolution structure of the human insulin receptor ectodomain: multi-modal inclusion of the insert domain. Structure 2016, 24, 469–476.

(16) Gutmann, T.; Kim, K. H.; Grzybek, M.; Walz, T.; Coskun, Ü. Visualization of ligand-induced transmembrane signaling in the full-length human insulin receptor. J. Cell Biol. 2018, 217, 1643–1649.

(17) Xu, Y.; Kong, G. K.-W.; Menting, J. G.; Margetts, M. B.; Delaine, C. A.; Jenkin, L. M.; Kiselyov, V. V.; De Meyts, P.; Forbes, B. E.; Lawrence, M. C. How ligand binds to the type 1 insulin-like growth factor receptor. Nat. Commun. 2018, 9, 821.

(18) Scapin, G.; Dandey, V. P.; Zhang, Z.; Prosise, W.; Hruza, A.; Kelly, T.; Mayhood, T.; Strickland, C.; Potter, C. S.; Carragher, B. Structure of the insulin receptor–insulin complex by single-particle cryo-EM analysis. Nature 2018, 556, 122–125.

(19) Li, J.; Choi, E.; Yu, H.; Bai, X.-c. Structural basis of the activation of type 1 insulin-like growth factor receptor. Nat. Commun. 2019, 10, 4567.

(20) Uchikawa, E.; Choi, E.; Shang, G.; Yu, H.; Bai, X.-c. Activation mechanism of the insulin receptor revealed by cryo-EM structure of the fully liganded receptor–ligand complex. eLife 2019, 8, e48630.

(21) Nielsen, J.; Brandt, J.; Boesen, T.; Hummelshøj, T.; Slaaby, R.; Schluckebier, G.; Nissen, P. Structural investigations of full-length insulin receptor dynamics and signalling. J. Mol. Biol. 2022, 434, 167458.

(22) Kavran, J. M.; McCabe, J. M.; Byrne, P. O.; Connacher, M. K.; Wang, Z.; Ramek, A.; Sarabipour, S.; Shan, Y.; Shaw, D. E.; Hristova, K.; Cole, P. A.; Leahy, D. J. How IGF-1 activates its receptor. eLife 2014, 3, e03772.

(23) Menting, J. G.; Lawrence, C. F.; Kong, G. K.-W.; Margetts, M. B.; Ward, C. W.; Lawrence, M. C. Structural congruency of ligand binding to the insulin and insulin/type 1 insulin-like growth factor hybrid receptors. Structure 2015, 23, 1271–1282.

(24) Lawrence, C. F.; Margetts, M. B.; Menting, J. G.; Smith, N. A.; Smith, B. J.; Ward, C. W.; Lawrence, M. C. Insulin mimetic peptide disrupts the primary binding site of the insulin receptor. J. Biol. Chem. 2016, 291, 15473–15481.

(25) Weis, F.; Menting, J. G.; Margetts, M. B.; Chan, S. J.; Xu, Y.; Tennagels, N.; Wohlfart, P.; Langer, T.; Müller, C. W.; Dreyer, M. K.; Lawrence, M. C. The signalling conformation of the insulin receptor ectodomain. Nat. Commun. 2018, 9, 4420.

(26) Gutmann, T.; Schäfer, I. B.; Poojari, C.; Brankatschk, B.; Vattulainen, I.; Strauss, M.; Coskun, Ü. Cryo-EM structure of the complete and ligand-saturated insulin receptor ectodomain. J. Cell Biol. 2019, 219, e201907210.

(27) Xiong, X. et al. A structurally minimized yet fully active insulin based on cone-snail venom insulin principles. Nat. Struct. Mol. Biol. 2020, 27, 615–624.

(28) Kirk, N. S.; Chen, Q.; Wu, Y. G.; Asante, A. L.; Hu, H.; Espinosa, J. F.; Martínez-Olid, F.; Margetts, M. B.; Mohammed, F. A.; Kiselyov, V. V.; Barrett, D. G.; Lawrence, M. C. Activation of the human insulin receptor by non-insulin-related peptides. Nat. Commun. 2022, 13, 5695.

(29) Kim, J.; Yunn, N.-O.; Park, M.; Kim, J.; Park, S.; Kim, Y.; Noh, J.; Ryu, S. H.; Cho, Y. Functional selectivity of insulin receptor revealed by aptamer-trapped receptor structures. Nat. Commun. 2022, 13, 6500.

(30) Park, J.; Li, J.; Mayer, J. P.; Ball, K. A.; Wu, J.; Hall, C.; Accili, D.; Stowell, M. H.; Bai, X.-c.; Choi, E. Activation of the insulin receptor by an insulin mimetic peptide. Nat. Commun. 2022, 13, 5594.

(31) Xiong, X. et al. Symmetric and asymmetric receptor conformation continuum induced by a new insulin. Nat. Chem. Biol 2022, 18, 511–519.

(32) Wu, M. et al. Functionally selective signaling and broad metabolic benefits by novel insulin receptor partial agonists. Nat. Commun. 2022, 13, 942.

(33) Li, J.; Park, J.; Mayer, J. P.; Webb, K. J.; Uchikawa, E.; Wu, J.; Liu, S.; Zhang, X.; Stowell, M. H.; Choi, E.; Bai, X.-c. Synergistic activation of the insulin receptor via two distinct sites. Nat. Struct. Mol. Biol. 2022, 29, 357–368.

(34) Xu, Y.; Margetts, M. B.; Venugopal, H.; Menting, J. G.; Kirk, N. S.; Croll, T. I.; Delaine, C.; Forbes, B. E.; Lawrence, M. C. How insulin-like growth factor I binds to a hybrid insulin receptor type 1 insulin-like growth factor receptor. Structure 2022, 30, 1098–1108.

(35) Choi, E.; Bai, X.-C. The activation mechanism of the insulin receptor: a structural perspective. Annu. Rev. Biochem. 2023, 92, 247–272.

(36) Bershatsky, Y. V. et al. Diversity of structural, dynamic, and environmental effects explain a distinctive functional role of transmembrane domains in the insulin receptor subfamily. Int. J. Mol. Sci. 2023, 24, 3906.

(37) Kuznetsov, A. S.; Zamaletdinov, M. F.; Bershatsky, Y. V.; Urban, A. S.; Bocharova, O. V.; Bennasroune, A.; Maurice, P.; Bocharov, E. V.; Efremov, R. G. Dimeric states of transmembrane domains of insulin and IGF-1R receptors: Structures and possible role in activation. Biochim. Biophys. Acta Biomembr. 2020, 1862, 183417.

(38) Frattali, A. L.; Treadway, J.; Pessin, J. Evidence supporting a passive role for the insulin receptor transmembrane domain in insulin-dependent signal transduction. J. Biol. Chem. 1991, 266, 9829–9834.

(39) Gardin, A.; Auzan, C.; Clauser, E.; Malherbe, T.; Aunis, D.; Crémel, G.; Hubert, P. Substitution of the insulin receptor transmembrane domain with that of glycophorin A inhibits insulin action. FASEB J. 1999, 13, 1347–1357.

(40) Longo, N.; Shuster, R.; Griffin, L. D.; Langley, S. D.; Elsas, L. J. Activation of insulin receptor signaling by a single amino acid substitution in the transmembrane domain. J. Biol. Chem. 1992, 267, 12416–12419.

(41) Whittaker, J.; Garcia, P.; Yu, G. Q.; Mynarcik, D. C. Transmembrane domain interactions are necessary for negative cooperativity of the insulin receptor. Mol. Endocrinol. 1994, 8, 1521–1527.

(42) Li, S.-C.; Deber, C. M.; Shoelson, S. E. An irregularity in the transmembrane domain helix correlates with the rate of insulin receptor internalization. Biochemistry 1994, 33, 14333–14338.

(43) Takahashi, K.; Yonezawa, K.; Nishimoto, I. Insulin-like growth factor I receptor activated by a transmembrane mutation. J. Biol. Chem. 1995, 270, 19041–19045.

(44) Hubbard, S. R.; Miller, W. T. Closing in on a mechanism for activation. eLife 2014, 3, e04909.

(45) Lee, J.; Miyazaki, M.; Romeo, G. R.; Shoelson, S. E. Insulin receptor activation with transmembrane domain ligands. J. Biol. Chem. 2014, 289, 19769–19777.

(46) De Meyts, P. Insulin/receptor binding: the last piece of the puzzle? What recent progress on the structure of the insulin/receptor complex tells us (or not) about negative cooperativity and activation. Bioessays 2015, 37, 389–397.

(47) Ward, C. W.; Menting, J. G.; Lawrence, M. C. The insulin receptor changes conformation in unforeseen ways on ligand binding: sharpening the picture of insulin receptor activation. Bioessays 2013, 35, 945–954.

(48) Mohammadiarani, H.; Vashisth, H. All-atom structural models of the transmembrane domains of insulin and type 1 insulin-like growth factor receptors. Front. Endocrinol. 2016, 7, 196560.

(49) Verma, J.; Vashisth, H. Molecular basis for differential recognition of an allosteric inhibitor by receptor tyrosine kinases. Proteins 2024, 1–18.

(50) Gorai, B.; Vashisth, H. Structural models of viral insulin-like peptides and their analogs. Proteins 2023, 91, 62–73.

(51) Gorai, B.; Vashisth, H. Progress in simulation studies of insulin structure and function. Front. Endocrinol. 2022, 13, 908724.

(52) Gorai, B.; Vashisth, H. Structures and interactions of insulin-like peptides from cone snail venom. Proteins 2022, 90, 680–690.

(53) Mohammadiarani, H.; Vashisth, H. Insulin mimetic peptide S371 folds into a helical structure. J. Comput. Chem. 2017, 38, 1158–1166.

(54) Vashisth, H. Flexibility in the insulin receptor ectodomain enables docking of insulin in crystallographic conformation observed in a hormone-bound microreceptor. Membranes 2014, 4, 730–746.

(55) Vashisth, H.; Abrams, C. F. All-atom structural models of insulin binding to the insulin receptor in the presence of a tandem hormone-binding element. Proteins 2013, 81, 1017–1030.

(56) Vashisth, H.; Maragliano, L.; Abrams, C. F. “DFG-flip” in the insulin receptor kinase is facilitated by a helical intermediate state of the activation loop. Biophys. J. 2012, 102, 1979–1987.

(57) Vashisth, H.; Abrams, C. F. Docking of insulin to a structurally equilibrated insulin receptor ectodomain. Proteins 2010, 78, 1531–1543.

(58) Arkhipov, A.; Shan, Y.; Das, R.; Endres, N. F.; Eastwood, M. P.; Wemmer, D. E.; Kuriyan, J.; Shaw, D. E. Architecture and membrane interactions of the EGF receptor. Cell 2013, 152, 557–569.

(59) Chavent, M.; Duncan, A. L.; Sansom, M. S. Molecular dynamics simulations of membrane proteins and their interactions: from nanoscale to mesoscale. Curr. Opin. Struct. Biol. 2016, 40, 8–16.

(60) Dominguez, L.; Foster, L.; Straub, J. E.; Thirumalai, D. Impact of membrane lipid composition on the structure and stability of the transmembrane domain of amyloid precursor protein. Proc. Natl. Acad. Sci. U. S. A. 2016, 113, E5281–E5287.

(61) Vanni, S.; Hirose, H.; Barelli, H.; Antonny, B.; Gautier, R. A sub-nanometre view of how membrane curvature and composition modulate lipid packing and protein recruitment. Nat. Commun. 2014, 5, 4916.

(62) Janosi, L.; Prakash, A.; Doxastakis, M. Lipid-modulated sequence-specific association of glycophorin A in membranes. Biophys. J. 2010, 99, 284–292.

(63) Hollingsworth, S. A.; Dror, R. O. Molecular dynamics simulation for all. Neuron 2018, 99, 1129–1143.

(64) Goossens, K.; De Winter, H. Molecular dynamics simulations of membrane proteins: An overview. J. Chem. Inf. Model. 2018, 58, 2193–2202.

(65) Marrink, S. J.; Corradi, V.; Souza, P. C.; Ingolfsson, H. I.; Tieleman, D. P.; Sansom, M. S. Computational modeling of realistic cell membranes. Chem. Rev. 2019, 119, 6184–6226.

(66) Li, P.-C.; Miyashita, N.; Im, W.; Ishido, S.; Sugita, Y. Multidimensional umbrella sampling and replica-exchange molecular dynamics simulations for structure prediction of transmembrane helix dimers. J. Comput. Chem. 2014, 35, 300–308.

(67) Dunton, T. A.; Goose, J. E.; Gavaghan, D. J.; Sansom, M. S.; Osborne, J. M. The free energy landscape of dimerization of a membrane protein, NanC. PLoS Comput. Biol. 2014, 10, e1003417.

(68) Cao, Y.; Yang, R.; Wang, W.; Jiang, S.; Yang, C.; Liu, N.; Dai, H.; Lee, I.; Meng, X.; Yuan, Z. Probing the formation, structure and free energy relationships of M protein dimers of SARS-CoV-2. Comput. Struct. Biotechnol. J. 2022, 20, 573–582.

(69) Bernardi, R. C.; Melo, M. C.; Schulten, K. Enhanced sampling techniques in molecular dynamics simulations of biological systems. Biochim. Biophys. Acta 2015, 1850, 872– 877.

(70) Yang, Y. I.; Shao, Q.; Zhang, J.; Yang, L.; Gao, Y. Q. Enhanced sampling in molecular dynamics. J. Chem. Phys. 2019, 151, 070902.

(71) Marrink, S. J.; Risselada, H. J.; Yefimov, S.; Tieleman, D. P.; De Vries, A. H. The MARTINI force field: coarse grained model for biomolecular simulations. J. Phys. Chem. B 2007, 111, 7812–7824.

(72) Kmiecik, S.; Gront, D.; Kolinski, M.; Wieteska, L.; Dawid, A. E.; Kolinski, A. Coarse-grained protein models and their applications. Chem. Rev. 2016, 116, 7898–7936.

(73) Gorai, B.; Sahoo, A. K.; Srivastava, A.; Dixit, N. M.; Maiti, P. K. Concerted interactions between multiple gp41 trimers and the target cell lipidome may be required for HIV-1 entry. J. Chem. Inf. Model. 2020, 61, 444–454.

(74) Prakash, A.; Janosi, L.; Doxastakis, M. Self-association of models of transmembrane domains of ErbB receptors in a lipid bilayer. Biophys. J. 2010, 99, 3657–3665.

(75) Prakash, A.; Janosi, L.; Doxastakis, M. GxxxG motifs, phenylalanine, and cholesterol guide the self-association of transmembrane domains of ErbB2 receptors. Biophys. J. 2011, 101, 1949–1958.

(76) Psachoulia, E.; Marshall, D.; Sansom, M. S. Molecular dynamics simulations of the dimerization of transmembrane *α*-helices. Acc. Chem. Res. 2010, 43, 388–396.

(77) De Jong, D. H.; Singh, G.; Bennett, W. D.; Arnarez, C.; Wassenaar, T. A.; Schafer, L. V.; Periole, X.; Tieleman, D. P.; Marrink, S. J. Improved parameters for the Martini coarse-grained protein force field. J. Chem. Theory Comput. 2013, 9, 687–697.

(78) Yang, Y.; Lee, M.; Fairn, G. D. Phospholipid subcellular localization and dynamics. J. Biol. Chem. 2018, 293, 6230–6240.

(79) Šali, A.; Blundell, T. L. Comparative protein modelling by satisfaction of spatial restraints. J. Mol. Biol. 1993, 234, 779–815.

(80) Martí-Renom, M. A.; Stuart, A. C.; Fiser, A.; Sánchez, R.; Melo, F.; Šali, A. Comparative protein structure modeling of genes and genomes. Annu. Rev. Biophys. Biomol. Struct. 2000, 29, 291–325.

(81) Shen, M.-y.; Šali, A. Statistical potential for assessment and prediction of protein structures. Protein Sci. 2006, 15, 2507–2524.

(82) Yesylevskyy, S. O.; Schäfer, L. V.; Sengupta, D.; Marrink, S. J. Polarizable water model for the coarse-grained MARTINI force field. PLoS Comput. Biol. 2010, 6, e1000810.

(83) Jo, S.; Kim, T.; Iyer, V. G.; Im, W. CHARMM-GUI: a web-based graphical user interface for CHARMM. J. Comput. Chem. 2008, 29, 1859–1865.

(84) Qi, Y.; Ingólfsson, H. I.; Cheng, X.; Lee, J.; Marrink, S. J.; Im, W. CHARMM-GUI Martini maker for coarse-grained simulations with the martini force field. J. Chem. Theory Comput. 2015, 11, 4486–4494.

(85) Abraham, M. J.; Murtola, T.; Schulz, R.; Páll, S.; Smith, J. C.; Hess, B.; Lindahl, E. GROMACS: High performance molecular simulations through multi-level parallelism from laptops to supercomputers. SoftwareX 2015, 1, 19–25.

(86) Humphrey, W.; Dalke, A.; Schulten, K. VMD: visual molecular dynamics. J. Mol. Graph. 1996, 14, 33–38.

(87) Wassenaar, T. A.; Pluhackova, K.; Bockmann, R. A.; Marrink, S. J.; Tieleman, D. P. Going backward: a flexible geometric approach to reverse transformation from coarse grained to atomistic models. J. Chem. Theory Comput. 2014, 10, 676–690.

(88) Lee, J.; Im, W. Transmembrane helix tilting: insights from calculating the potential of mean force. Phys. Rev. Lett. 2008, 100, 018103.

(89) Daura, X.; Gademann, K.; Jaun, B.; Seebach, D.; Van Gunsteren, W. F.; Mark, A. E. Peptide folding: when simulation meets experiment. Angew. Chem. Int. Ed. 1999, 38, 236–240.

(90) Chothia, C.; Levitt, M.; Richardson, D. Helix to helix packing in proteins. J. Mol. Biol. 1981, 145, 215–250.

(91) Enkavi, G.; Javanainen, M.; Kulig, W.; Róg, T.; Vattulainen, I. Multiscale simulations of biological membranes: the challenge to understand biological phenomena in a living substance. Chem. Rev. 2019, 119, 5607–5774.

(92) von Heijne, G. Proline kinks in transmembrane *α*-helices. J. Mol. Biol. 1991, 218, 499–503.

(93) Dong, H.; Sharma, M.; Zhou, H.-X.; Cross, T. A. Glycines: role in *α*-helical membrane protein structures and a potential indicator of native conformation. Biochemistry 2012, 51, 4779–4789.

(94) Wilman, H. R.; Shi, J.; Deane, C. M. Helix kinks are equally prevalent in soluble and membrane proteins. Proteins 2014, 82, 1960–1970.

(95) Cabail, M. Z.; Li, S.; Lemmon, E.; Bowen, M. E.; Hubbard, S. R.; Miller, W. T. The insulin and IGF1 receptor kinase domains are functional dimers in the activated state. Nature Commun. 2015, 6, 6406.

(96) Xu, Y.; Kirk, N. S.; Venugopal, H.; Margetts, M. B.; Croll, T. I.; Sandow, J. J.; Webb, A. I.; Delaine, C. A.; Forbes, B. E.; Lawrence, M. C. How IGF-II binds to the human type 1 insulin-like growth factor receptor. Structure 2020, 28, 786–798.

