## Supporting Information for "Spontaneous Dimerization and Distinct Packing Modes of Transmembrane Domains in Receptor Tyrosine Kinases"

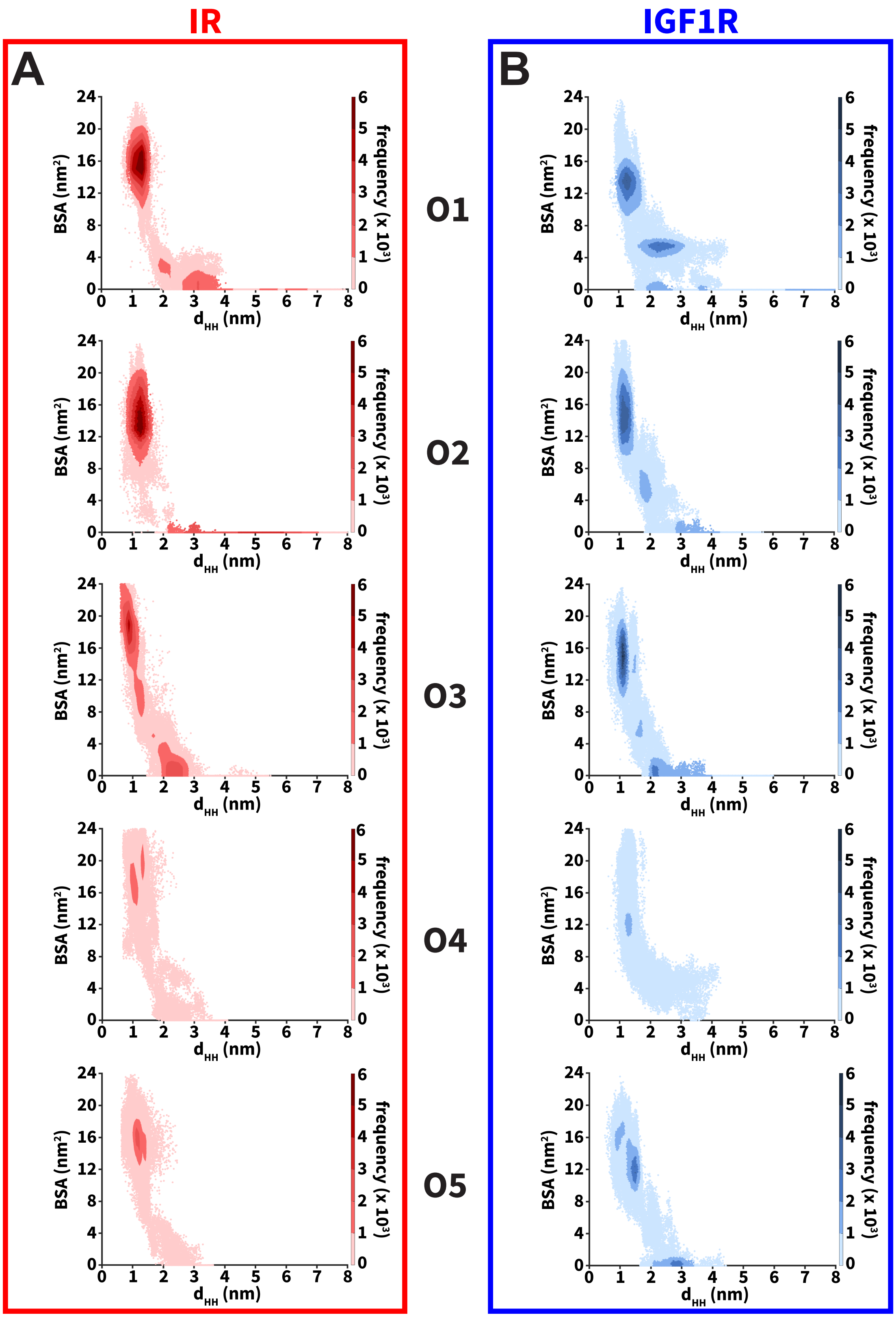


**Figure S1.** Buried surface area (BSA) distributions as a function of the interhelical distance ($d_{HH}$) computed from all CG simulations for each initial orientation (labeled O1 through O5) of (A) IR and (B) IGF1R TMDs.

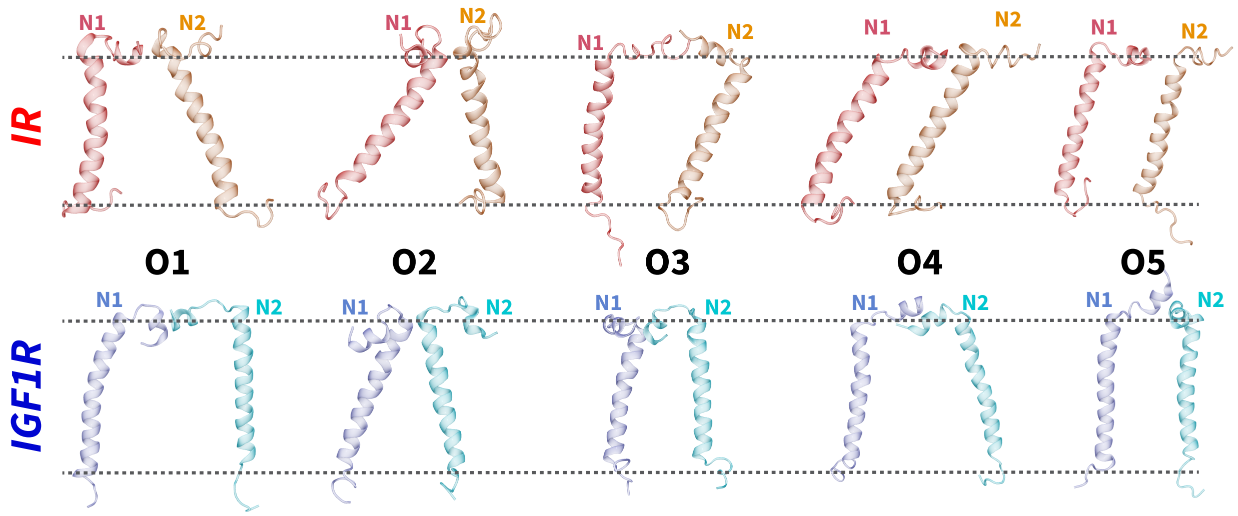


**Figure S2.** Cartoon representations highlighting the transient association of TMDs through interactions between the N-terminus of each TMD (labeled N1 and N2): IR TMDs (*top*) and IGF1R TMDs (*bottom*). The horizontal dotted gray lines indicate the approximate location of the lipid bilayer.

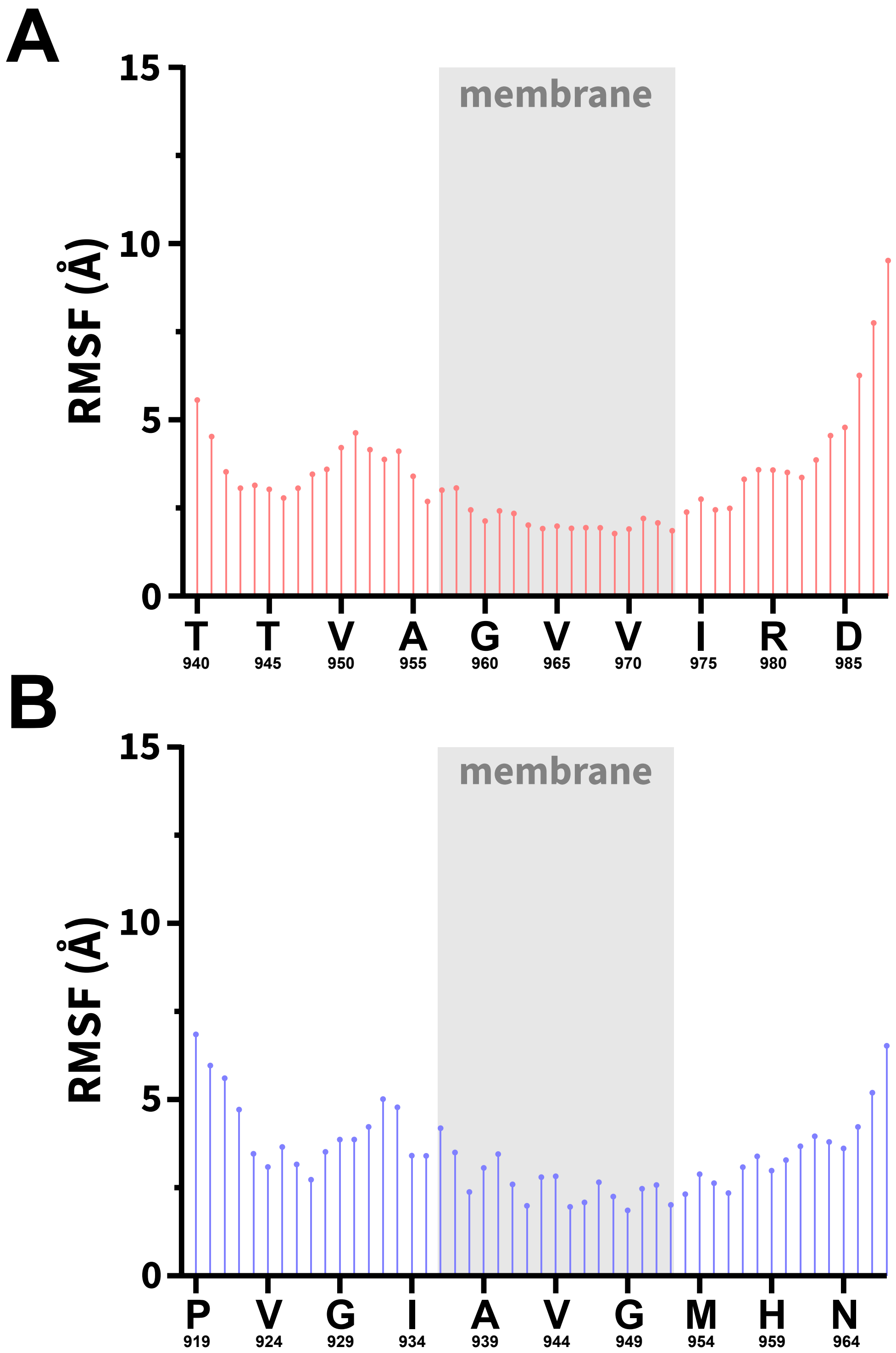


**Figure S3.** The RMSF per residue computed from all CG simulations of (A) IR TMDs and (B) IGF1R TMDs. The gray rectangle marks residues embedded in the membrane.

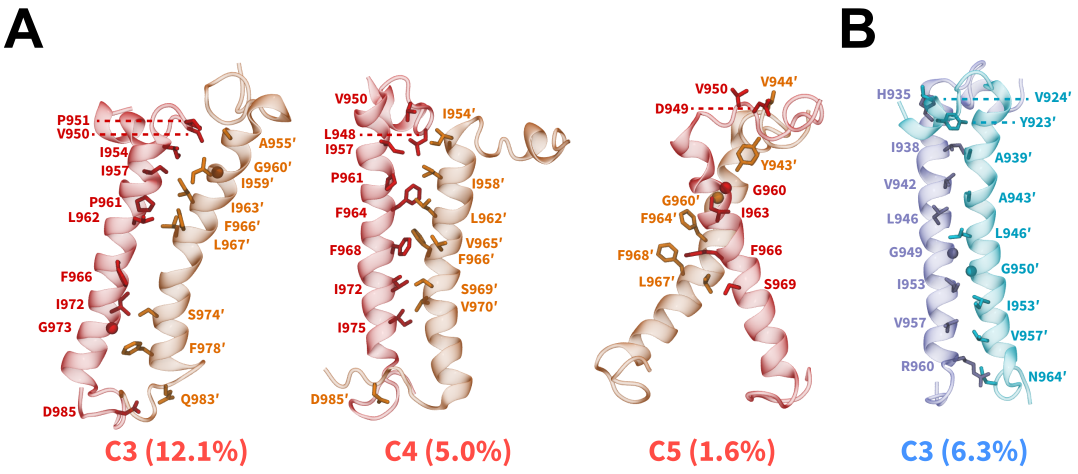


**Figure S4.** Cartoon representations highlighting the dimeric states of TMDs derived from smaller-sized conformational clusters of (A) IR and (B) IGF1R TMDs. The names of conformational clusters (C3, C4, and C5) and their sizes (%) are labeled. See also Figure 4.

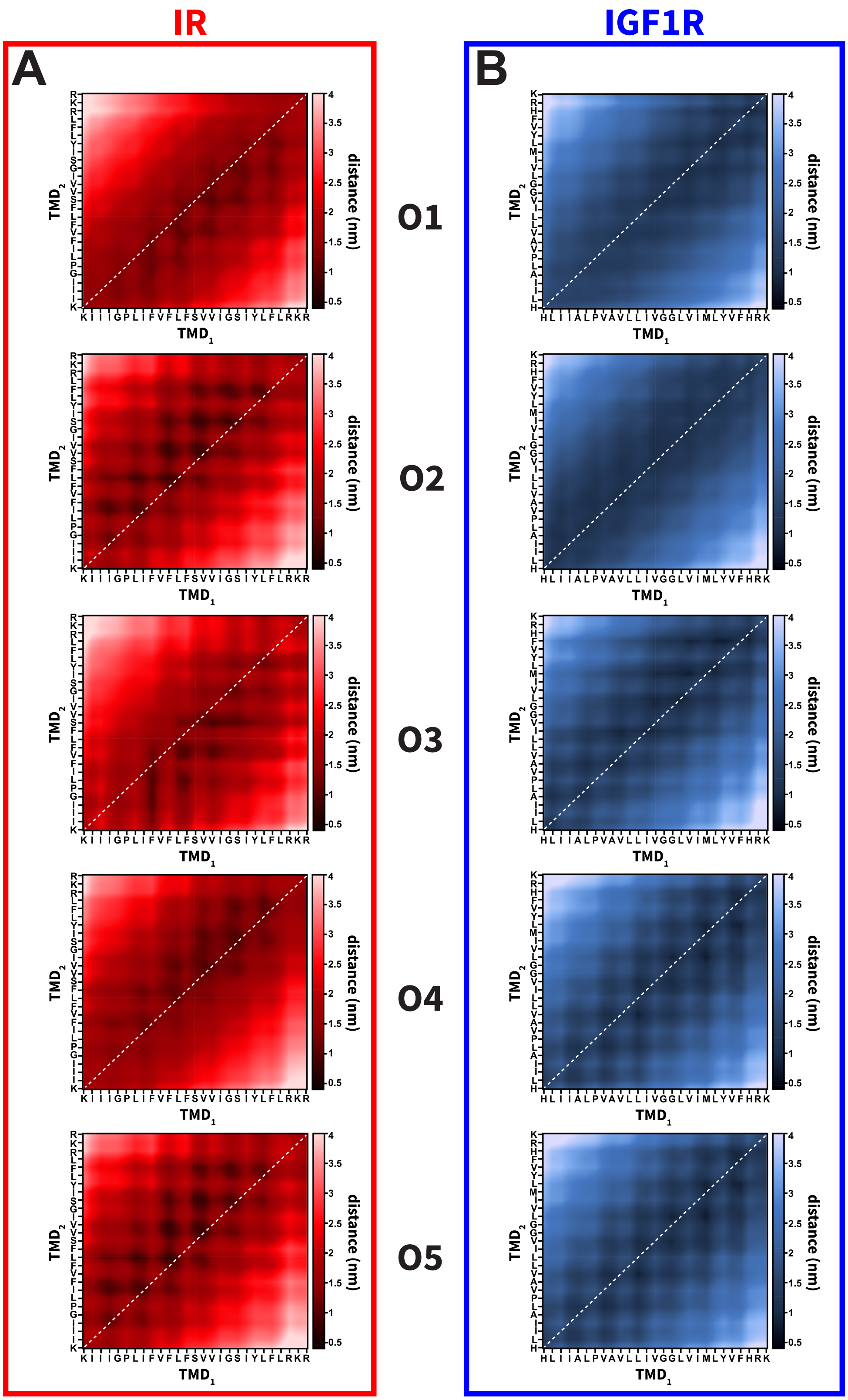


**Figure S5.** The averaged distance contact maps computed from all CG simulations of (A) IR TMD and (B) IGF1R TMD dimers. The dotted white line denotes the symmetry axis in each contact map.

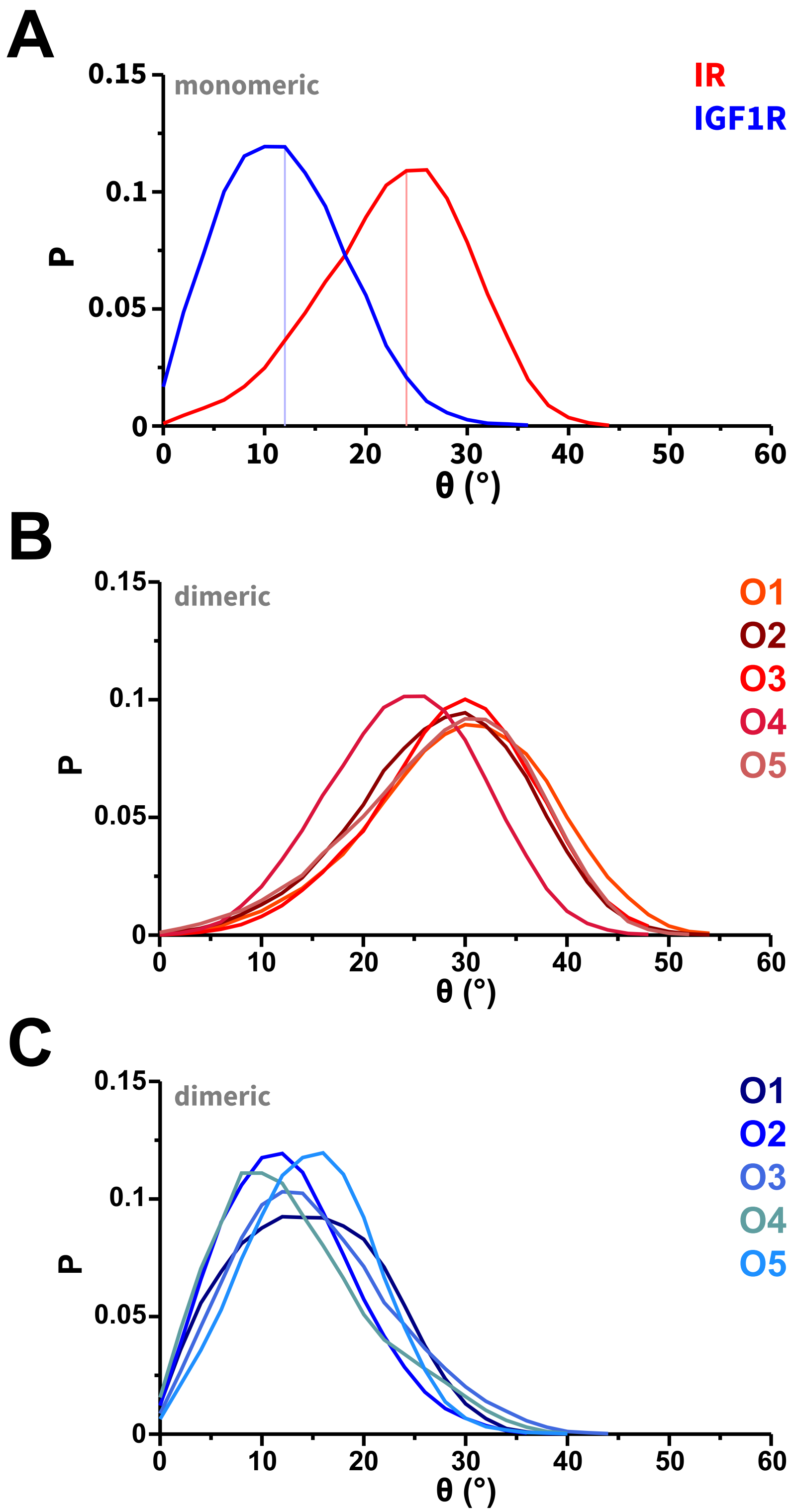


**Figure S6.** The probability distributions of the tilt-angle (θ) computed from CG simulations of: (A) monomeric IR/IGF1R TMDs and (B, C) dimeric IR/IGF1R TMDs. Distributions are colored uniquely (red, IR; blue, IGF1R) to mark the specific distribution derived from CG simulations based on different initial orientations (labeled O1 through O5).

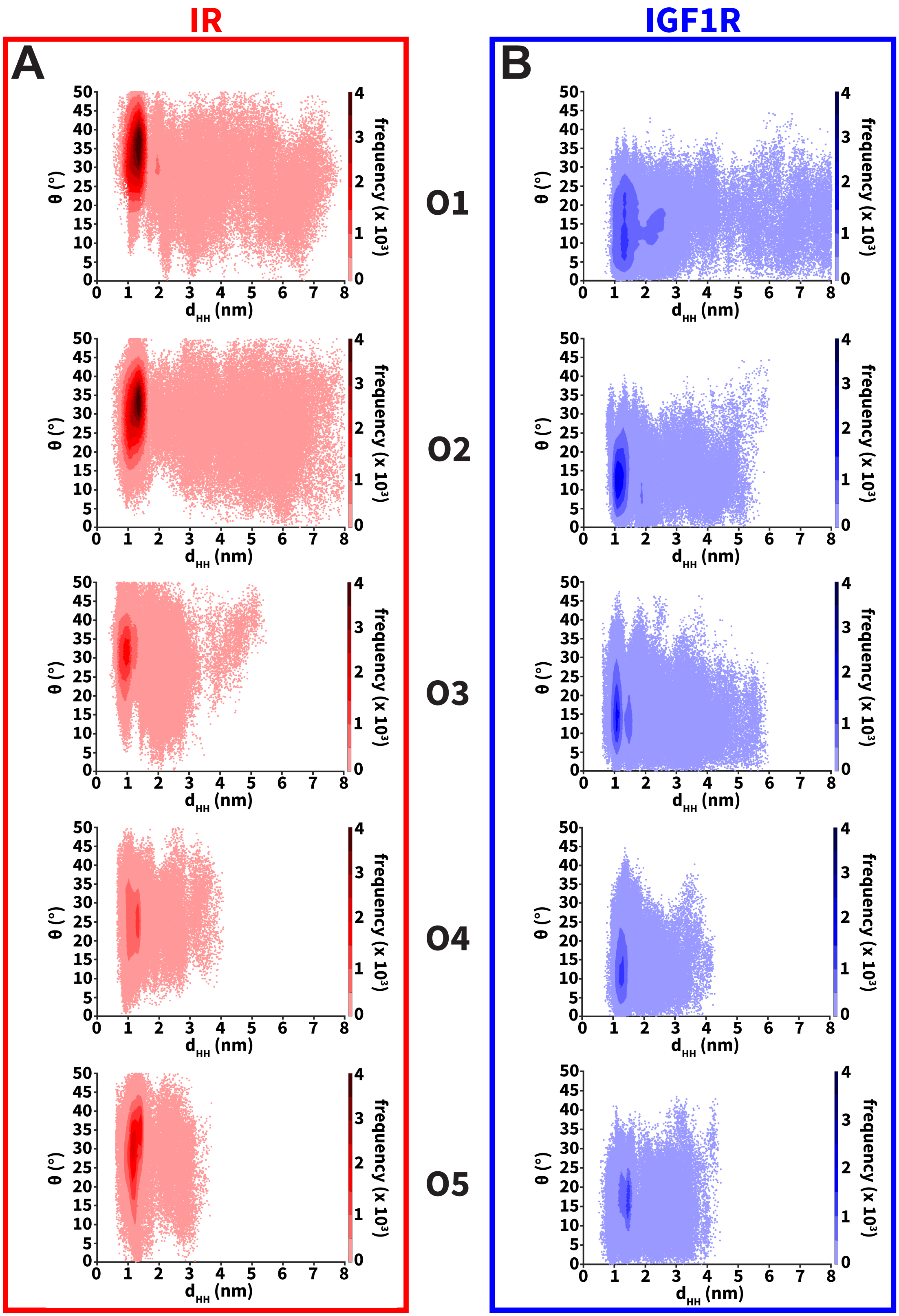


**Figure S7.** Distributions of θ vs. $d_{HH}$ computed from CG simulations of (A) IR TMDs and (B) IGF1R TMDs for each initial orientation (labeled O1 through O5).

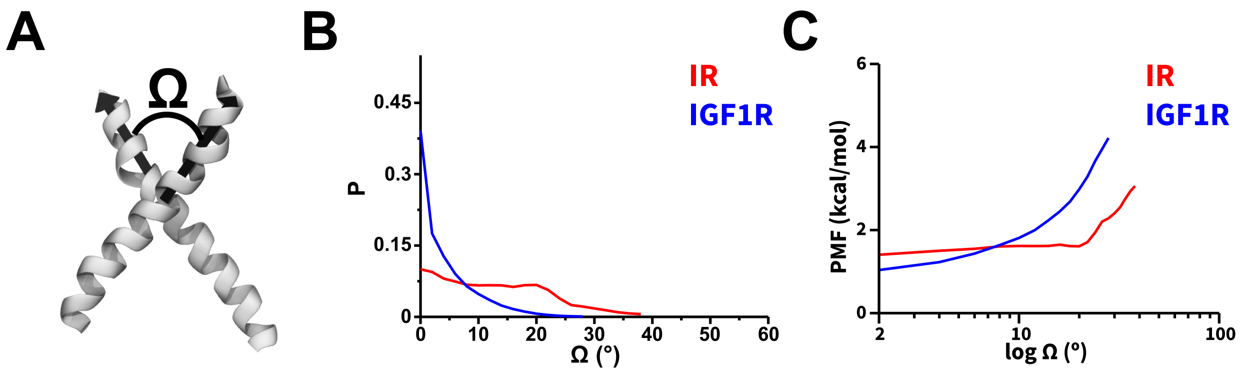


**Figure S8.** (A) A schematic highlighting the vectors along TMD helices that are used to define the crossing-angle (Ω), (B) the probability distributions of Ω, and (C) the potential of mean force (PMF) highlighting the free energy change (kcal/mol) as a function of Ω computed from all CG simulations of IR (red) and IGF1R (blue) TMDs.

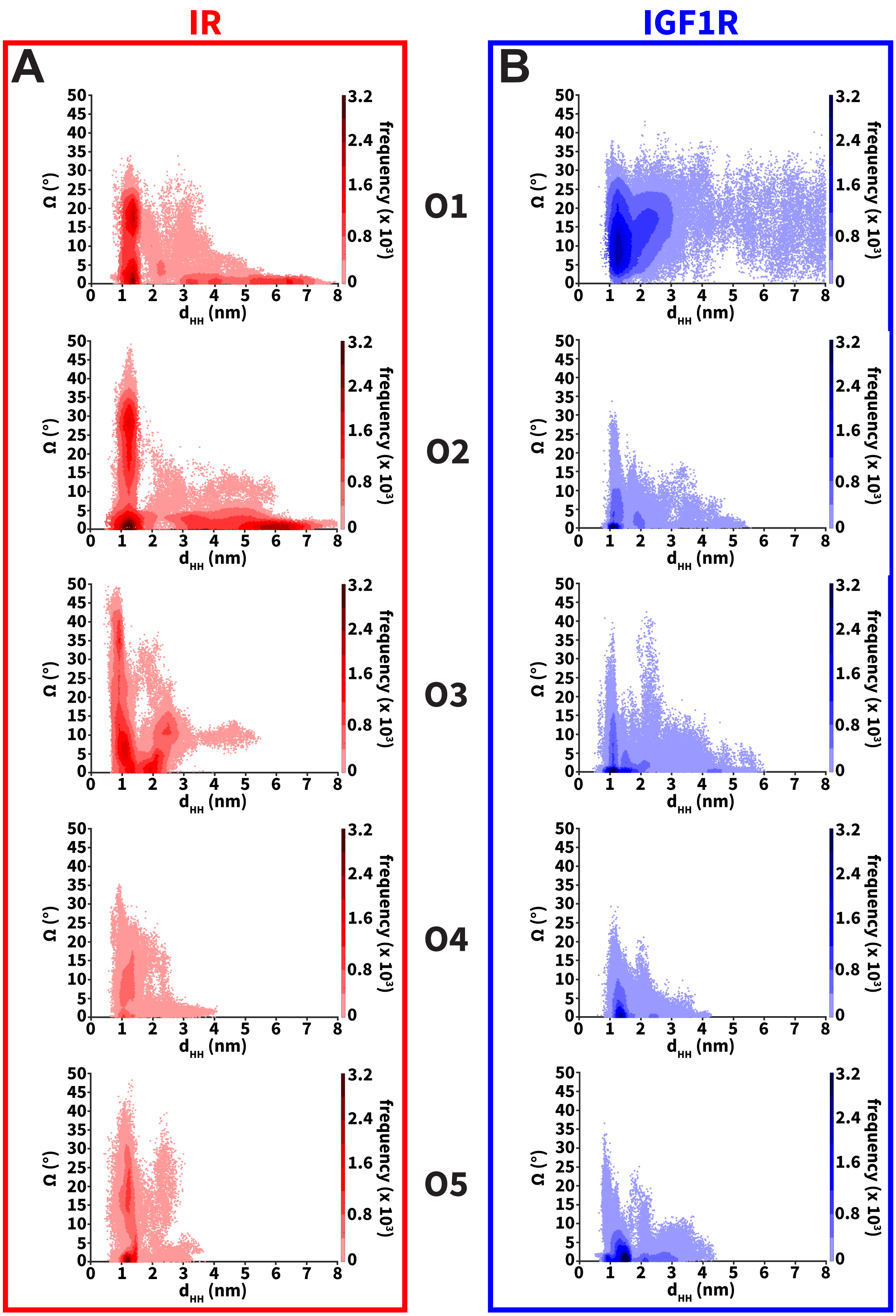


**Figure S9.** Distributions of Ω vs. $d_{HH}$ computed from CG simulations of (A) IR and (B) IGF1R TMDs for each initial orientation (labeled O1 through O5).
